# Structure-Activity-Relationship Advancement, Animal Validation, and Target Engagement of a Novel Assembly Modulator Small Molecule Chemical Series with Pan-Cancer Selective Cytotoxicity

**DOI:** 10.1101/2025.02.28.640839

**Authors:** Anuradha F. Lingappa, Suguna Mallesh, Maya Michon, Devaraju Shivarudraswamy, Jim Lin, M. Dharma Prasad, Debendranath Dey, Dennis Solas, Bingyu B. Li, Andreas Mueller-Schiffmann, David W. Andrews, Carsten Korth, Stevan Pecic, Vishwanath R. Lingappa

## Abstract

PAV-620, PAV-621, and PAV-805 are chemical analogs from a protein assembly modulating small molecule lead series that displays broad anti-cancer activity. These compounds are cytotoxic to all cancers in the NCI-60 cancer cell line screen, which is representative of diverse blood, brain, breast, colon, lung, ovarian, prostate, renal, and skin cancers, while nontoxic to healthy human peripheral blood mononuclear cells at doses up to 20uM. The compounds are as efficacious as Paclitaxel in reducing tumor growth and metastasis in the aggressive 4T1 mouse allograft model for triple negative breast cancer. The compounds’ mechanism of action in primary carcinoma cells under conditions inducing programmed cell death was found to be distinct from the cytotoxic mechanisms of Staurosporine, Paclitaxel, and Etoposide. The safest compound from the series thus far identified, PAV-805, is nontoxic to mice following twice-daily doses of 10 mg/kg. PAV-805 was shown to target a subset of protein disulfide isomerase (PDI) in a dynamic multi-protein complex enriched in the cancer hallmark proteins involved in reprogramming energy metabolism. These data suggest a new approach to cancer therapeutics, with selectivity arising not from targeting PDI itself, an abundant cellular protein, but from targeting a particular disease-associated assembly state of PDI as a therapeutically exploitable layer of cancer biology.

## Introduction

We recently described PAV-951, a novel chemical compound originally identified as a modulator of HIV capsid assembly in a cell free protein synthesis and assembly (CFPSA) system.^1^ Cell-free systems have played an important role in the history of biochemistry and molecular cell biology. Such systems were used in the 19^th^ century to demonstrate that fermentation of sugar to alcohol is possible outside living cells, in the 1960’s to decipher the genetic code, and in the 1970’s to establish the signal hypothesis of protein targeting.^2–4^ We used CFPSA to translate viral capsid mRNA and reconstitute capsid formation from those nascent chains, as an approximation to what happens in infected cells.^5,6^ Formation of capsids indistinguishable from the authentic capsid observed in infected cells by all criteria (biochemical, biophysical, and electron microscopic appearance) was achieved and demonstrated to occur via an energy-dependent, host-catalyzed pathway.^5–8^ We then adapted the CFPSA assay into a phenotypic drug screen to identify assembly-modulating small molecules^1,5,7–11^. Hit compounds identified from the CFPSA screen have demonstrated anti-viral activity via disruption of critical protein-protein interactions at the host-viral interface.^8–10,12^

The anti-viral activity of PAV-951 was validated against infectious HIV in cell culture studies and the compound was then counter-screened against cancer.^1^ While unorthodox, our pursuit of anti-viral compounds for cancer therapeutics has a compelling logic. A vast body of literature connects viruses with cancer ^13–15^ Oncogenic viral proteins facilitate viral propagation and predispose cells to tumorigenesis by various mechanisms, including the disruption or modification of biochemical pathways which are critical for maintaining homeostasis, cell cycle control, and apoptosis.^15,16^

Human Papilloma Virus (HPV) is a well-studied example of an oncogenic virus that demonstrates this phenomenon.^15,17^ The HPV E7 viral protein is itself oncogenic— expression of HPV E7 in a transgenic mouse is sufficient to trigger cervical cancer development.^15,17–19^ HPV E7 viral proteins have not been observed to display any intrinsic enzymatic activities, but they are known to associate with and modify the functions of cellular protein complexes.^15,20,21^ For example, HPV E7 induces metabolic reprogramming in the host cells by directly binding Hypoxia-Inducible Factor-1a (HIF-1a).^22,23^ Such metabolic reprogramming, is a strategy employed by multiple diverse families of viruses in hijacking a host cell. ^16,24–31^ However, it is also one of the classic hallmarks of cancer.^32,33^. The connection between the viral modification of host multi-protein complexes and the subcellular changes which may lead to cancer development, formed the basis for our hypothesis that *<u>compounds which normalize protein-protein interactions at the host-viral interface might also restore the diverse homeostatic controls that are disrupted in neoplastic cells.</u>*^1^

When tested against tumor cell lines and in mouse models of cancer, the anti-viral assembly-modulating compound PAV-951 was observed to have remarkably broad activity against cancer.^1^ Anti-cancer activity was observed against 80 of 80 different tumor cell lines (including all 60 of the NCI-60 screen) and significantly inhibited tumor growth in two mouse xenograft studies on lung and colon cancer.

The potential of PAV-951 to be further developed into a cancer therapeutic is limited by risk of toxicity (PAV-951 is only nontoxic to mice at doses at or below 2.5 mg/kg) and variability within the broad anti-cancer activity (while it displayed activity against all cancers tested, its efficacy against some cancer cell lines were an order of magnitude less potent than against others).^1^ We sought to overcome these liabilities and optimize the chemistry by assessment of new chemical analogs of PAV-951.

We published an initial structure-activity-relationship (SAR) dataset on PAV-951 and three of its analogs, PAV-436, PAV-541, and PAV-621.^1^ In the initial SAR set, we showed that small changes to the chemical structure of PAV-951 could significantly improve or completely eliminate the anti-cancer activity observed in cell culture.^1^ Here, we share data on PAV-621 and other chemical analogs from the PAV-951 series, including PAV-620 and PAV-805, which demonstrate superior safety while improving upon the breadth and potency of the anti-cancer activity. Cellular and biochemical methods are used to further elucidate the functional target and develop testable hypotheses on the compounds’ mechanism of action. The studies presented here support the hypothesis that the PAV-951 lead series targets an allosteric site controlling metabolic reprogramming across cancers.

## Results

### Structure-Activity-Relationship (SAR) Exploration of Assembly Modulating Chemical Series Identifies Compounds with Increased Breadth of Anti-Cancer Efficacy And Broad Therapeutic Window

Chemical analogs of PAV-951 were synthesized and assessed in the National Cancer Institute 60 cell line screen (NCI-60). Compounds were tested against up to 60 tumor cell lines representative of brain cancer, breast cancer, lung cancer, colon cancer, leukemia, melanoma, ovarian cancer, renal cancer, and prostate cancer. SAR exploration resulted in the identification of compounds with strikingly improved efficacy relative to PAV-951. *See* **Figure 1** for NCI-60 data on PAV-951 and 15 of its chemical analogs at a single dose of 2.5uM, and **Figure S1** for NCI-60 data from five-dose titrations of two lead compounds, PAV-620 and PAV-805.

**Figure 1.**
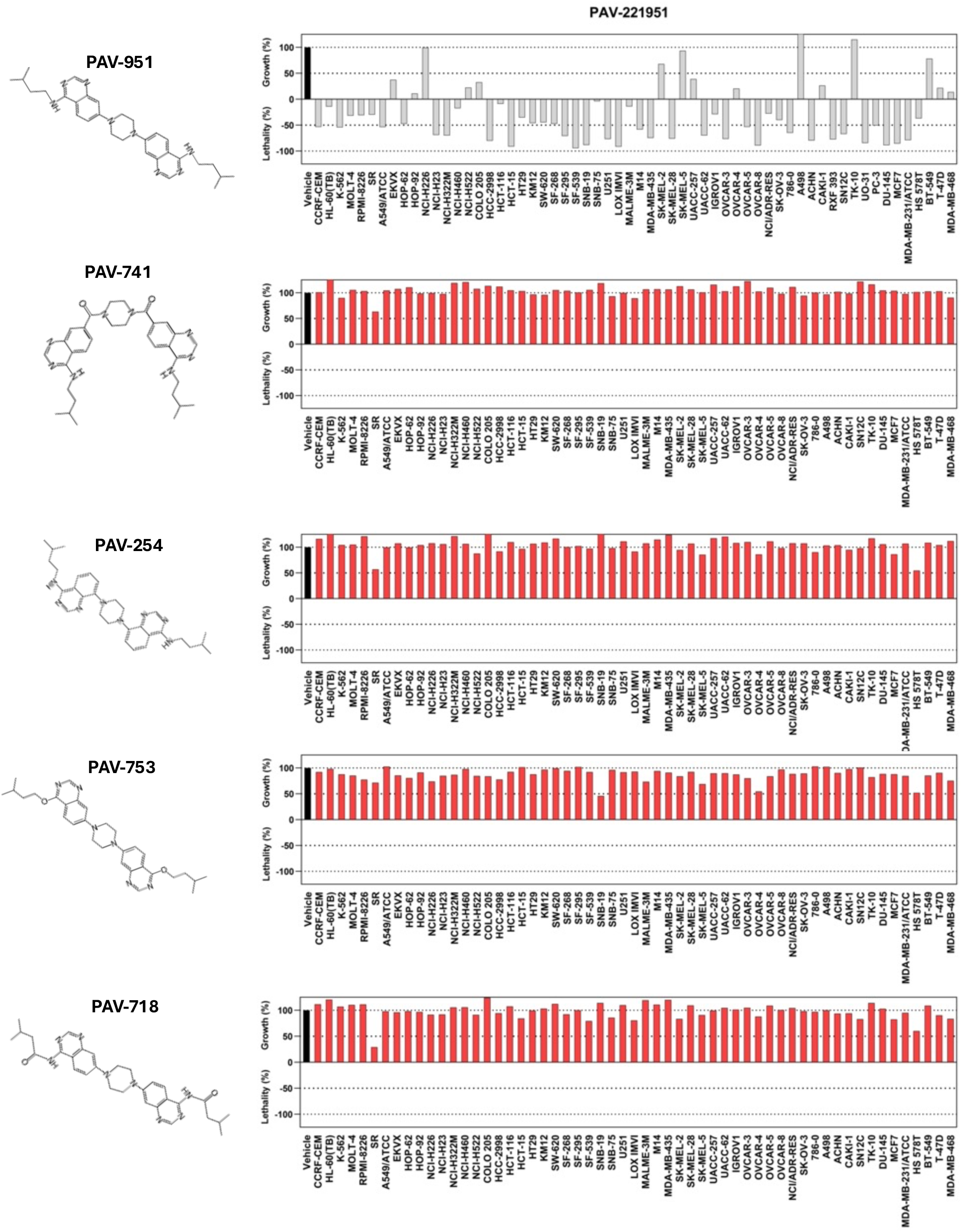
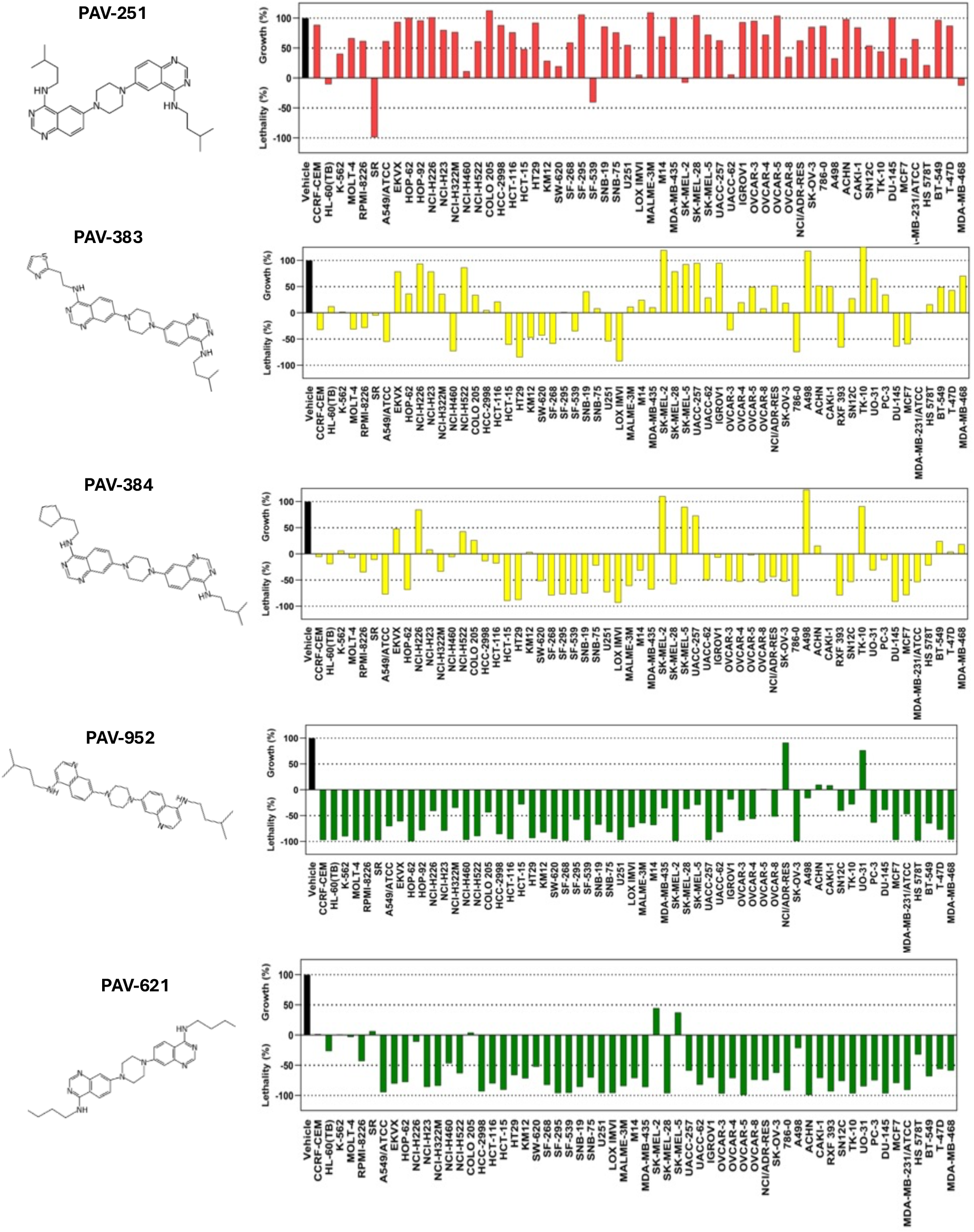
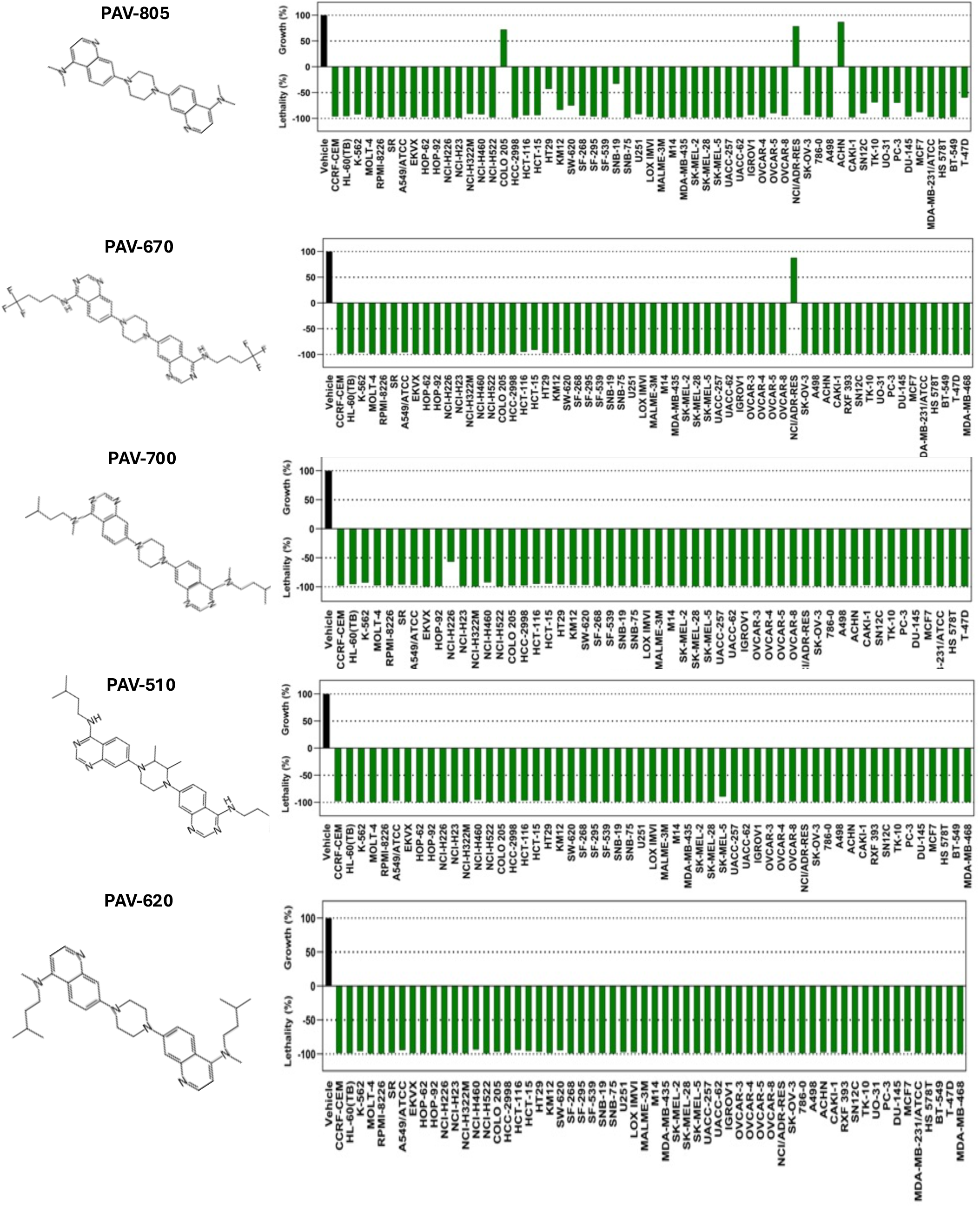
Assessment of PAV-951 and its analogs in the NCI-60 Screen. **Figure 1** shows the chemical structures of PAV-951 and 15 of its chemical analogs alongside activity of each compound against the NCI-60 panel at a single dose of 2.5uM. Each bar represents a different tumor cell line and cell viability of each cell line upon drug treatment is shown relative to the vehicle treated control and the number of cells at time 0. Values between 0 and 100 represent percent growth inhibition. Values between 0 and -100 represent percent cellular lethality. Graphs are color coded where red indicates loss, green indicates gain, and yellow indicates no substantial change in the anti-cancer activity of analogs compared to PAV-951.

Following the NCI-60 results assessing the compounds at 2.5uM, we conducted dose titrations to assess the cytotoxicity of active compounds in A549 lung cancer and HT29 colon cancer cells at 20uM, 5uM, 1.25uM, and 0.325uM. *See* **Figures 2A** and **2B**.

**Figure 2.**
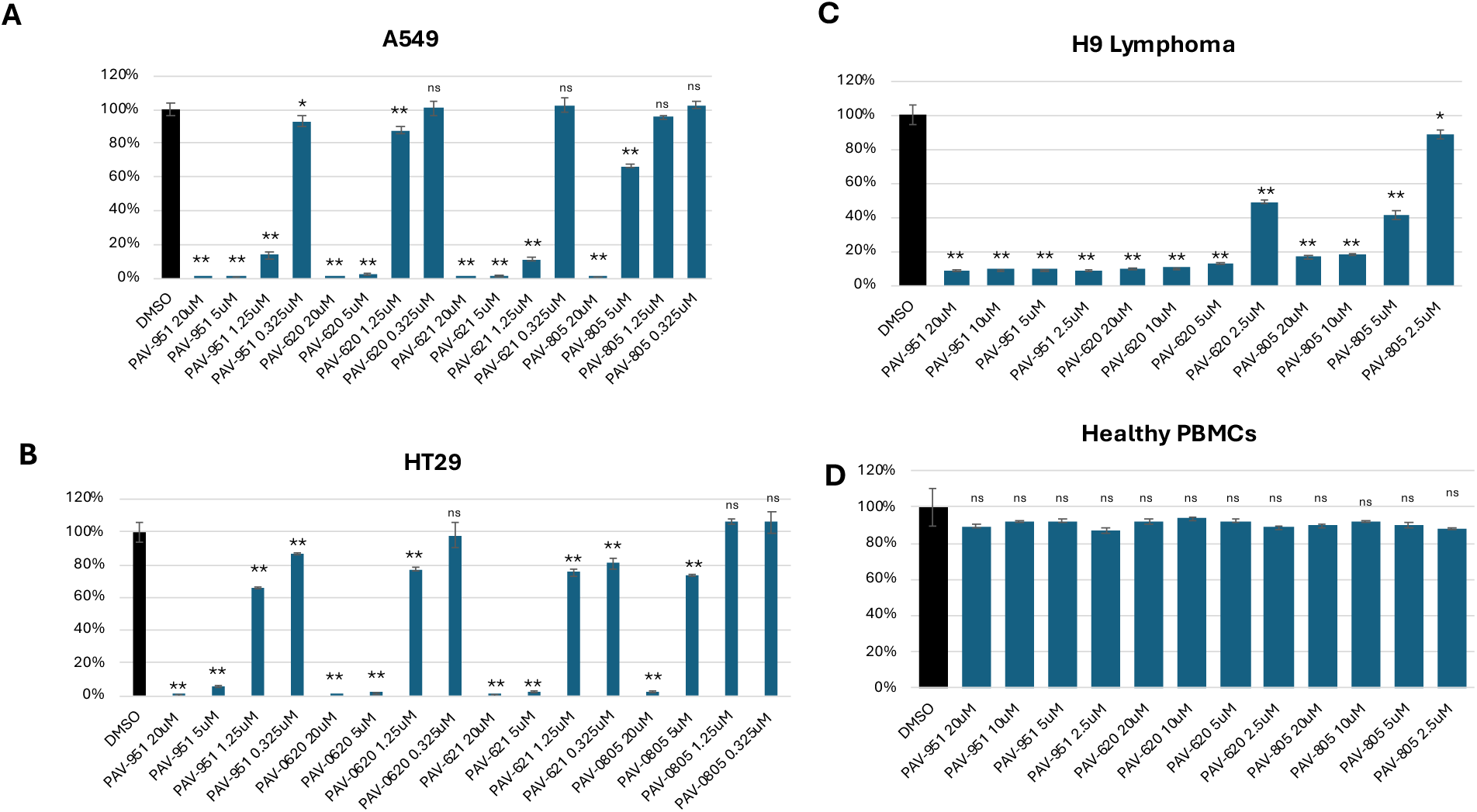
Cell Viability in A549, HT29, H9, and PBMC Cells Following Dose-Titrations of Assembly Modulator Compounds. A549, HT29, H9 and PBMC cells were seeded at a density of 5,000 cells/well. A549 and HT29 cells were incubated at 37oC for 24 hours and then treated with either DMSO or compound at 20uM, 5uM, 1.25uM, or 0.325uM. The treated plates were incubated for 72 hours then cell viability was measured using a Titer-Glo Assay and readings were graphed as percent of control. See **Figures 2A** and **2B**. H9 and PBMC cells were treated at the time of plating with either DMSO or compound at 20uM, 10uM, 5uM, or 2.5uM. The cells were incubated at 37oC for 72 hours then cell viability was measured using an AlamarBlue^TM^ assay and readings were graphed as percent of control. See **Figures 2C** and **2D**. Error bars indicate standard deviation between triplicate-repeated conditions. Statistical analysis was conducted between the DMSO and compound-treated cells using a t-test and statistical significance is indicated with asteriks. The difference in cell viability between DMSO and compound-treated cells was found to be significant (p-value < 0.000005) for every H9 testing condition examined. The difference in cell viability between DMSO and compound treated cells was found to be significant for only one PBMC condition (2.5uM of PAV-951, p-value = 0.02).

To characterize the therapeutic window between safety and efficacy, compounds were then assessed against H9 lymphoma cells versus peripheral blood mononuclear cells (PBMCs) from healthy individuals under matched conditions. Compounds were found to be non-toxic to PBMCs, while maintaining cytotoxicity against the lymphoma. *See* **Figures 2C** and **2D**. Thus, SAR exploration identified multiple advanced compounds that display increased cytotoxicity to tumor cell lines, relative to the original compound PAV-951 while maintaining a broad therapeutic window between efficacy and safety.

### Validation of the Compounds’ Anti-Cancer Activities in the 4T1 Mouse Allograft Model for Metastatic Triple Negative Breast Cancer at Nontoxic Doses

PAV-621 was the first analog of PAV-951 identified with significantly improved activity. Therefore, we sought to assess the anti-cancer activity of PAV-621 in an animal model of cancer. In advance of an efficacy study, pharmacokinetics (PK) and multi-day tolerability was profiled in mice. PAV-621 was found to be safe in mice following daily intraperitoneal (IP) doses of 2.5 mg/kg for 14 days. See Table S1.

4T1 is an aggressive triple-negative mouse breast cancer cell line which spontaneously metastasizes in mice.^36,37^ After 4T1 cells are implanted in mice, a palpable tumor forms within a week and progresses to a tumor volume of > 2,000 mm3 within 30 days.^36,37^ An advantage of using this allograft model, in contrast to the xenograft studies which were used to assess PAV-951, is that the mice are immunocompetent and the interface between a test compound and the immune system is preserved.

In a first allograft study, 4T1 tumors were implanted into mammary fat pads of immunocompetent BALB/c mice which were then treated with vehicle, Paclitaxel (Taxol), or PAV-621. Tumor volume was measured overtime and, after 28 days, animals were euthanized and the number of metastatic lung nodules was counted. When compound was administered at the time of tumor implantation, Paclitaxel inhibited tumor growth by 36% and reduced metastasis by 42%, and PAV-621 inhibited tumor growth by 47% and reduced metastasis by 39%, relative to control.

In one treatment group (referred to as the “late” treatment group), animals were administered vehicle until the tumors had grown to a volume of 50-60mm^3^, at which point the tumor was established and representative of a stage 2 presentation. PAV-621 was then administered daily to the late treatment group for the rest of the study. Between day 7 (when treatment began) and day 28, PAV-621 inhibited tumor growth by 46% and reduced metastasis by 32%. *See* **Figures 3A** and **3B** for efficacy data and **Figure S3** for animal body weights during the study. Throughout the study, no adverse clinical signs were observed. No significant variations in hematology were observed in compound-treated animals. See Figure S4.

**Figure 3.**
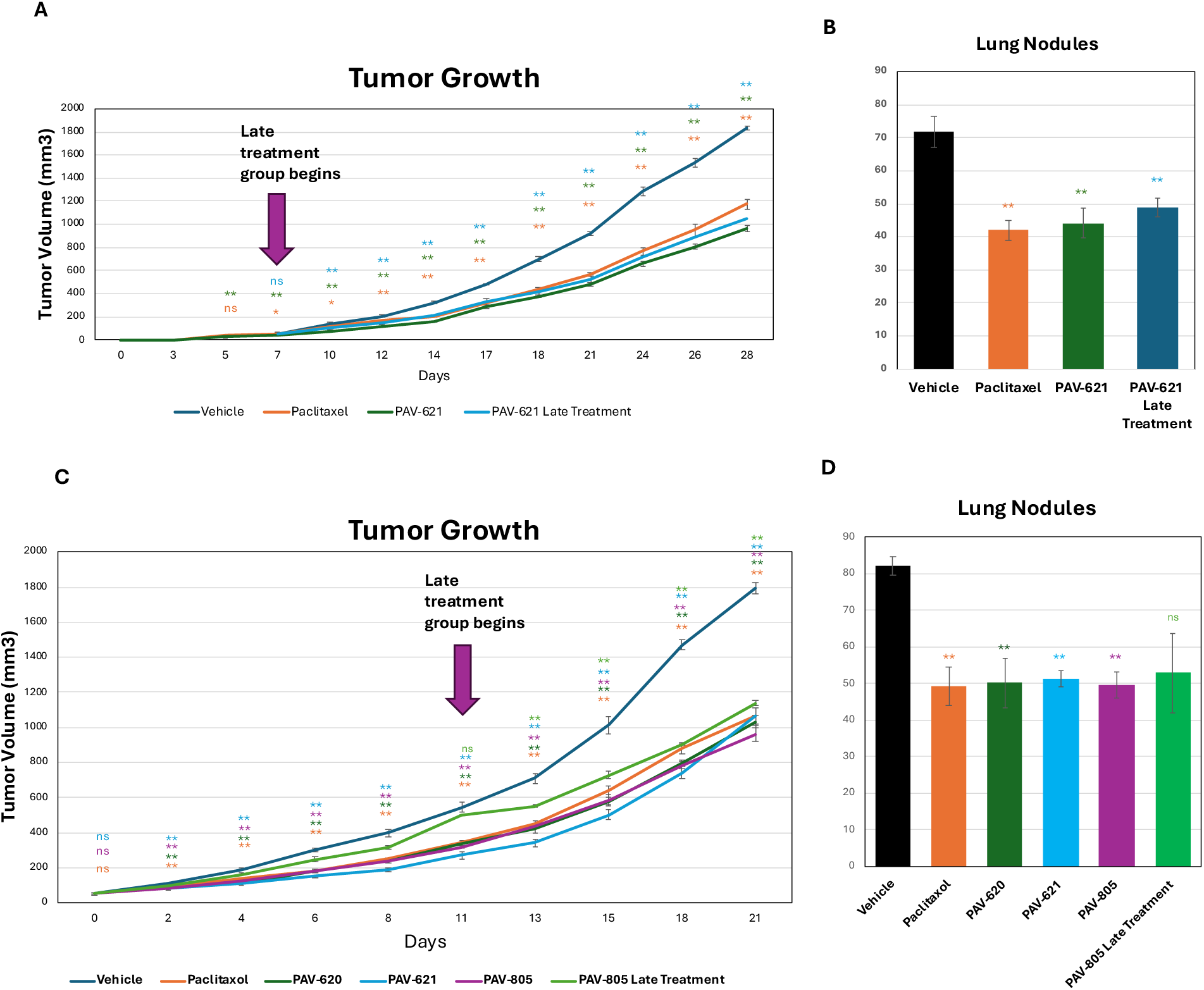
Activity of PAV-620 and PAV-805 in the 4T1 allograft model. 4T1 mouse breast cancer tumors were transplanted onto mice and treated with compound. **Figures 3A** and **3B** show the results of the first allograft study where vehicle, Paclitaxel, or PAV-621 were administred at the time of tumor implantation or beginning when the tumor reached a size of 50-60mm^3^ (late treatment). **Figures 3C** and **3D** show the results of the second allograft study where vehicle, Paclitaxel, PAV-620, PAV-621, or PAV-805 were administered beginning when the tumor reached a size of 50mm^3^ or 500mm^3^ (late treatment). In the first allograft study, PAV-621 was administered at a dose of 2 mg/kg daily. In the second allograft study, PAV-621 was administered at a dose of 2.5 mg/kg daily and PAV-805 was administered at a dose of 10 mg/kg daily. PAV-620 was administered at 5 mg/kg of compound daily for the first week of the study. However, on day 8, due to concerns about toxicity, the dose of PAV-620 was reduced to 1.5 mg/kg daily for the remainder of the study. In both studies, animals in the vehicle-only group were dosed daily and animals in the Paclitaxel group received compound at a dose of 2 mg/kg, administered every-other-day. Upon completion of the study, animals were euthanized and the number of metastatic cancer cell colonies in the lung tissue were counted. **Figure 3A** and **C** shows tumor growth over time and **Figure 3B** and **D** shows the number of lung nodules at euthanasia, for each treatment group where results have been averaged and error bars indicate standard deviation. Statistical analysis was performed using a t-test comparing vehicle and compound treated groups for each day and statistical significance is indicated with asterisk. Significant inhibition of tumor growth was observed as early as day two of the study and remained significant through day 21. On day 21, the p-values for tumor growth inhibition of the Paclitaxel, PAV-620, early PAV-805, and late PAV-805 treatment groups were found to be 1.05×10^-7^, 1.08×10^-8^, 1.4×10^-8^, and 2.1×10^-6^, respectively. On day 21, reduction of lung metastasis for the Paclitaxel, PAV-620, early PAV-805, and late treatment groups were found to be 0.0002, 0.001, 0.00002, and .077 respectively.

While the first allograft study was ongoing, we identified two compounds from the series, PAV-620 and PAV-805, that appeared substantially safer in mice. Both PAV-620 and PAV-805 could be administered to mice at single IP dose of 10 mg/kg without any observed symptoms of toxicity. When assessed for multi-dose safety, compounds were well-tolerated in mice. No symptoms of toxicity were observed in male or female mice that received twice-daily administration of PAV-805 at 10 mg/kg, for 14 days. *See* Table S1. When administered to mice at an IP dose of 5 mg/kg, PAV-620, PAV-621, and PAV-805 displayed similar exposure in the plasma over 24 hours. *See* Table S2.

In a second allograft study, 4T1 tumors were implanted into mammary fat pads of immunocompetent BALB/c mice which were then treated with vehicle, Paclitaxel, PAV-620, PAV-621, or PAV-805. Tumor volume was measured over time and, after 21 days, animals were euthanized and the number of metastatic lung nodules was counted. In the second allograft study compound administration in the early treatment groups began when the tumors were at a volume of 50mm^3^. Under those conditions, Paclitaxel inhibited tumor growth by 42% and reduced metastasis by 40%, PAV-620 inhibited tumor growth by 44% and reduced metastasis by 39%, PAV-621 inhibited tumor growth by 52% and reduced metastasis by 38%, and PAV-805 inhibited tumor growth by 48% and reduced metastasis by 40%, relative to control. *See* **Figures 3C and 3D** for efficacy and **Figure S5** for body weight over the study.

In the second allograft study, a “late” treatment group received received vehicle daily until day 11 when the tumors had grown to a volume of 500mm^3^, at which point the cancer was in the exponential growth phase and already metastatic, representative of a stage 3 or 4 presentation. PAV-805 was then administered for the rest of the study. In the late treatment group, PAV-805 showed an immediate effect. Between day 11 (when PAV-805 treatment began) and day 13, PAV-805 inhibited tumor growth by 68.9%. Between day 11 and day 21 (when the study was completed), PAV-805 treatment inhibited tumor growth by 48.9%. As a point of comparison for the other treatment conditions, between day 0 and day 21 (including the time when animals only received vehicle), late administration of PAV-805 inhibited tumor growth by 38% and reduced metastasis by 30%, relative to control. *See* **Figures 3C and 3D for efficacy and Figure S5** for body weight over the study.

Thus, PAV-620, PAV-621, and PAV-805 display potent anti-cancer activity in the 4T1 mouse allograft model as measured by both inhibition of the primary tumor and reduction of metastasis. The compounds’ effect on tumor growth inhibition appears to be equally potent on early and late stage presentations. The anti-cancer activity of PAV-621 was reproducible across two different allograft studies. The three analogs show comparable activity against the cancer *in vivo*, despite having substantially different mouse safety profiles.

### PAV-620 and PAV-805 Exhibit a Unique Mechanism of Action in Primary Carcinomas

As the compounds from the PAV-951 series showed comparable efficacy to Paclitaxel in the 4T1 model, we then sought to compare the mechanism-of-action of our compounds and other cytotoxic agents in metastatic breast cancer cells. We selected PAV-805 as the main compound to use for mechanistic and target-identification studies due to its increased mouse safety, following the rationale that any observed cytotoxicity to cancer cells would be target-specific rather than off-target “collateral damage”.

Cells from two primary breast carcinomas (BB5 and BB7) were seeded on non-adherent plates and treated either with DMSO, Staurosporine, Etoposide, Paclitaxel, PAV-620, or PAV-805. Live cell stains ChromaLIVE snd AnnexinV were added during seeding (to assess cell morphology and apoptosis, respectively). After 24 hours, was added to stain DNA. Images of the cells were captured and analyzed for intensity, morphology, and texture features that were then analyzed in Python/R Under these conditions, cells would normally undergo anoikis, a specialized form of programmed cell death that is triggered when healthy, anchorage-dependent cells detach from their extracellular matrix.^38,39^ A Random Forest Classifier (RFC) program sampled images of DMSO and Staurosporine-treated cells to train on distinguishing between control and dead cells. The RFC program then determined the percentage of dead cells in plates treated with increasing doses of Paclitaxel, Etoposide, PAV-620, and PAV-805. At the concentrations and times tested here, Paclitaxel and Etoposide did not increase cell death beyond the ongoing anoikis in cells from either carcinoma. *See* **Figures 4A** and **4B**. PAV-620 and PAV-805 greatly increased cell death in the BB7 carcinoma cells and slightly increased cell death in the BB5 carcinoma cells. *See* **Figures 4A** and **4B**.

**Figure 4.**
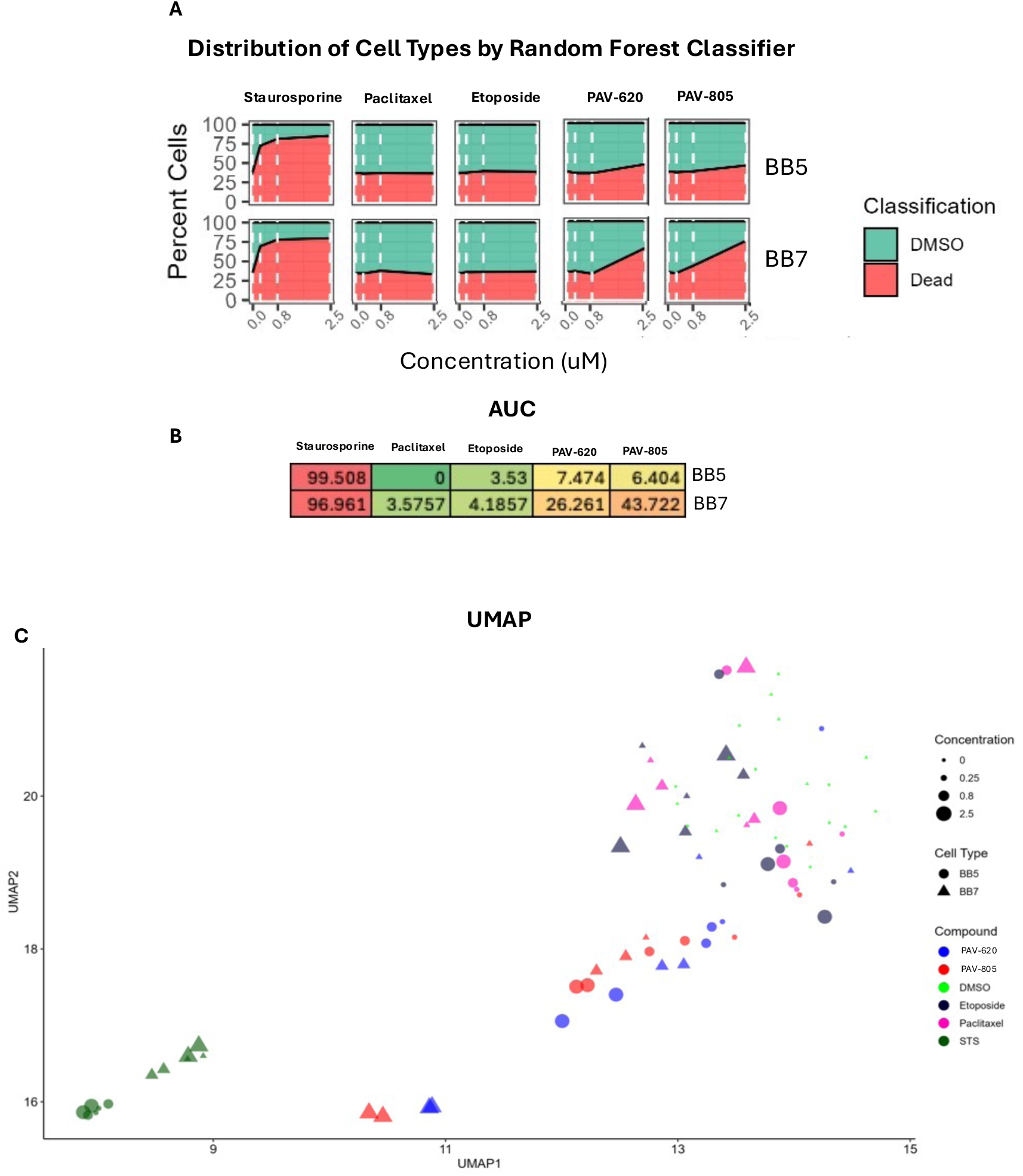
Assessment of Staurosporine, Paclitaxel, Etoposide, PAV-620 and PAV-805 in primary carcinoma. BB5 and BB7 carcinoma cells were seeded on non-adherent plates at 2,000 cells/well with live cell stains and treated with DMSO, 0.25uM, 0.8uM, or 2.5uM Staurosporine, Etoposide, Paclitaxel, PAV-620, or PAV-805 on day 0. After 24 hours, Hoechst was added and cells were imaged. Results were analyzed in Python/R. **Figures 4A** and **4B** indicate percentage of dead cells and the corresponding area under the curve (normalized against DMSO) for each treatment condition. **Figure 4C** shows a 2-dimensional UMAP analysis.

A 2-dimensional UMAP analysis was run using the images of stained cells. PAV-620 and PAV-805-treated BB5 and BB7 clustered together, in a distinct region of the UMAP from DMSO, Staurosporine, Paclitaxel, or Etoposide-treated cells. *See* **Figure 4C**. Furthermore, as the concentrations of PAV-620 and PAV-805 increased (size of the symbols increases), the symbols representing cell images moved progressively further away from the less affected cells.

### PAV-951 and PAV-805 Target Overlapping but Distinct Sets of Proteins

PAV-951, the parent compound from the cancer assembly modulator series, was determined to target a multi-protein complex by energy-dependent drug resin affinity chromatography (eDRAC)^1^ PAV-805 was coupled to a resin from the same attachment point as PAV-951. *See* **Figure S6** for structures of the resins**. e**DRAC was conducted on the PAV-951 and PAV-805 resins in parallel using A549 lung cancer cell extract as starting material. Analysis of the eluates by silver stain and western blot showed the two compounds target overlapping, but distinct, sets of proteins. *See* **Figures 5A-F** for quantitation of western blot bands and **Figures S7A** and **S7B** for pictures of the silver stains and western blots.

**Figure 5.**
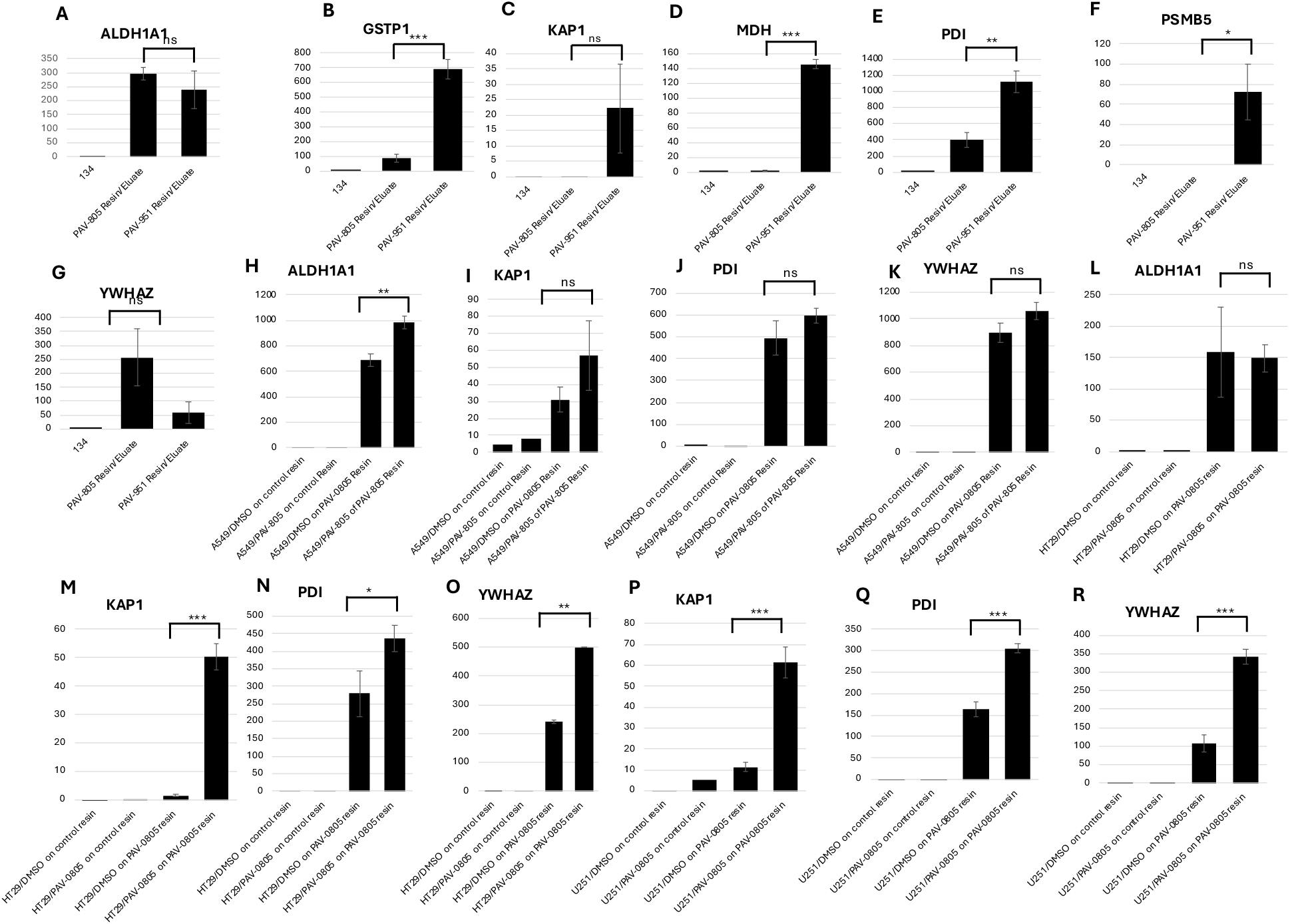
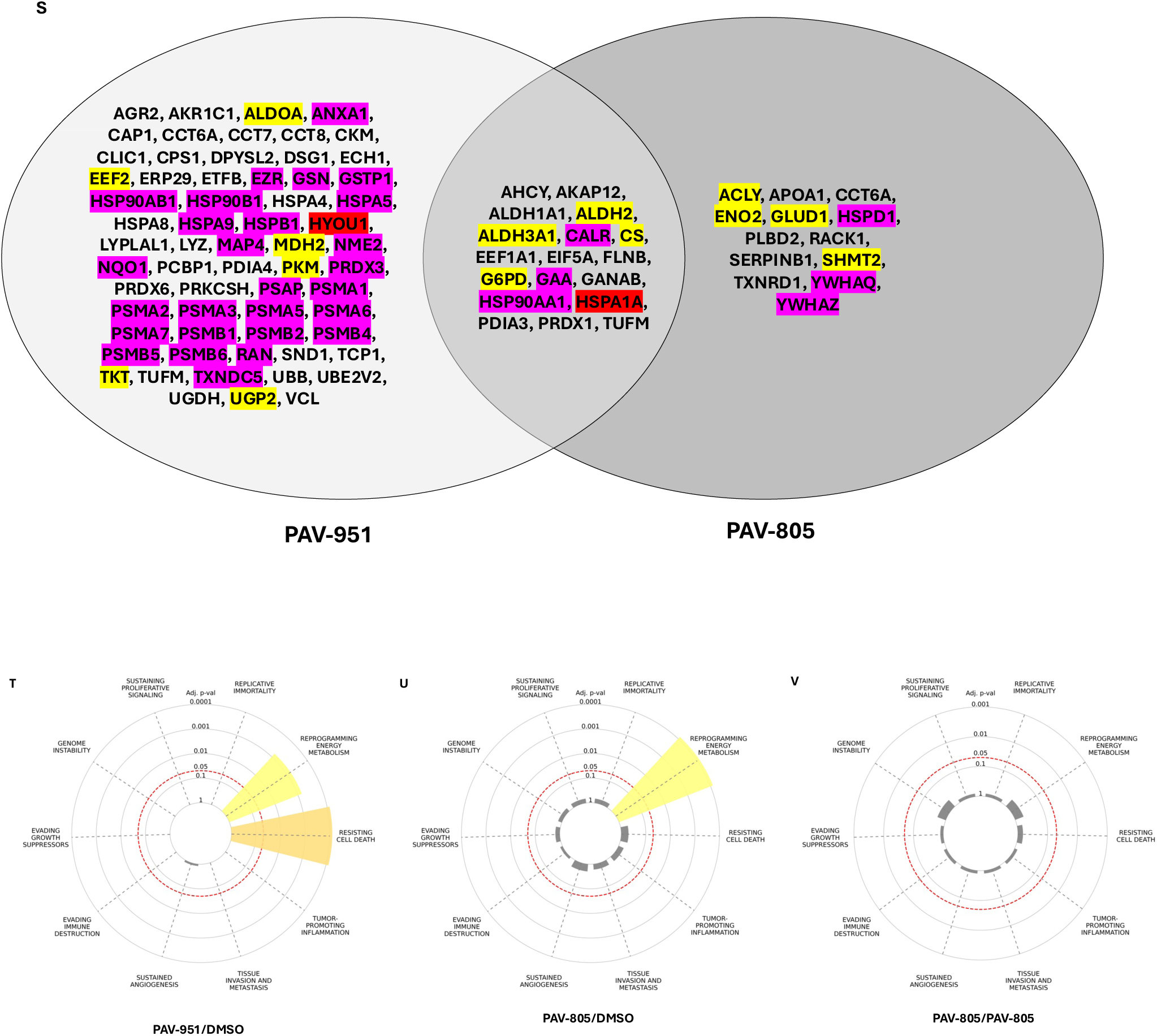
Drug Resin Affinity Chromatography with PAV-951 and PAV-805 Resins. A549, HT29, and U251 cells were treated with DMSO or PAV-805 and lysate was prepared and used for eDRAC, repeated in triplicate on the PAV-951 and PAV-805 resins, as well as in single-point on a control resin. The eDRAC eluates were run on SDS PAGE and analyzed by western blot and MSMS. **Figures 5A-F** show quantitation of western blot bands in eDRAC experiments comparing the eluate composition from DMSO-treated A549 samples run on the PAV-951 and PAV-805 resins in parallel. **Figures 5G-J** show quantitation of western blot bands in experiments comparing the eluate composition from DMSO versus PAV-805 treated A549 samples run on the PAV-805 resin. **Figures 5K-N** show quantitation of western blot bands in experiments comparing the eluate composition from DMSO versus PAV-805 treated HT29 samples run on the PAV-805 resin. **Figures 5O-Q** show quantitation of western blot bands in experiments comparing the eluate composition from DMSO versus PAV-805 treated U251 samples run on the PAV-805 resin. For the western blot quantitation, graphs show the average density of the protein band detected in triplicate-repeated samples and error bars indicate standard deviation. Statistical analysis for experiments that were run in parallel was done using a t-test and statistical significance is denoted by asterisk. Where multiple bands were present on the western blot for a given protein, quantitation shows the 100kDa KAP1 band, the 57kDa PDI band, and the 25kDa MDH band. **Figure 5R** shows a Venn Diagram of gene products identified by MSMS in the PAV-951 and PAV-805 resin eluates, grouped by the proteins only identified in the PAV-951 target, proteins identified in both the PAV-951 and PAV-0805 targets, and proteins only identified in the PAV-0805 target. The lists of proteins comprising each of the eluate components were searched in a database for genes associated with the hallmarks of cancer.^32,40^. Association with a particular cancer hallmark is based on the enrichment of hallmark-associated genes identified relative to a reference set.^40^ Color coding indicates whether a gene product is associated with a cancer hallmark of resisting cell death (pink), reprogramming energy metabolism (yellow), or both (red). **Figures 5S-U**, adapted from Menyhart et. al., show association of the PAV-951 and PAV-805 resin eluate components. The set of proteins in the DMSO-treated PAV-951 resin eluate is significantly enriched for the hallmarks of reprogramming energy metabolism and resisting cell death. The set of proteins in the DMSO-treated PAV-805 resin eluate is significantly enriched for the hallmark of reprogramming energy metabolism. The set of proteins in the PAV-805-treated PAV-805 resin eluate is not enriched for any hallmarks of cancer. For the analysis of MSMS data used in **Figures 5R-U**, proteins identified in the negative control resin eluates were excluded and proteins not detected in all three of the triplicate-repeated samples were excluded.

The PAV-805 resin eluates were sent for tandem mass spectrometry (MSMS) analysis. A set of 181 proteins were detected by MSMS in the DMSO-treated PAV-805 resin eluate *See* **Table S3**. Of these, 115 proteins were not detected in eluate collected from a negative control resin under matched conditions *See* **Table S3**. Of these, 30 proteins (ACLY, AHCY, AKAP12, ALDH1A1, ALDH2, ALDH3A1, APOA1, CALR, CCT6A, CS, EEF1A1, E1F5A, ENO2, FLNB, G6PD, GAA, GLUD1, HSP90AA1, HSPA1A, HSPD1, PDIA3, PLBD2, PRDX1, RACK1, SERPINB1, SHMT2, TUFM, TXNRD1, YWHAQ, and YWHAZ) were detected in all three triplicate-repeat samples. *See* **Table S3.**

The set of proteins comprising the PAV-805 target was compared to set of proteins previously-identified as comprising the PAV-951 target^1^. The MSMS data corroborated the western blot findings that the eluates contain overlapping, but distinct sets of proteins.

### The Composition of the PAV-805 Target Changes with Compound Treatment in Multiple Cancer Cell Types

Given the observed broad activity on diverse cancers in cell culture, we wanted to characterize the PAV-805 target in diverse cancer cell types. eDRAC was conducted on the PAV-805 resin using extract from A549 lung cancer, HT29 colon cancer, and U251 brain cancer cells. Western blot analysis identified KAP1, PDI, and YWHAZ in the resin eluates from all three starting materials and ALDH1A1 in the resin eluate from A549 and HT29, but not U251 cells. *See* **Figures 5G-Q** for quantitation of protein bands and **Figure S7C** for pictures of western blots.

eDRAC experiments were conducted in parallel using extract from cells treated with DMSO or PAV-805 for 20 hours prior to lysis. Western blot analysis identified significantly more protein in the eluates from PAV-805-treated cells compared to the DMSO-treated cells. *See* **Figures 5G-Q** for quantitation of protein bands and **Figure S7C** for pictures of western blots. The majority of KAP1 detected by western blot was a fragment at ∼50kDa. *See* **Figure S7C** for pictures of western blots. However, for all three cancer cell types, Western blot analysis showed the appearance of a KAP1 band at 100kDa in the eluates from cells treated with PAV-805. *See* **Figures 5H**, **5L, and 5O** for quantitation of the 100kDa KAP1 band and **Figure S7C** for pictures of western blots.

The PAV-805 resin eluates from PAV-805-treated A549 cell extracts were analyzed by MSMS. A set of 134 proteins were detected by MSMS in the resin eluate from PAV-805-treated cells. *See* **Table S3**. Of these, 43 proteins were not detected in eluate collected from a negative control resin under matched conditions. *See* **Table S3**. Of these, 8 proteins (AHCY, ALDH1A1, ALDH3A1, CKM, HSPA5, AHNAK, AMY2A, and PDIA3) were detected in all three triplicate-repeat samples. *See* Table S3. Of these, all but two (AHNAK and AMY2A) were also detected by MSMS in the triplicate-repeat samples of DMSO-treated eluates (although a small amount of HSPA5 and CKM had been detected in the negative control resin eluates for the DMSO treated set and were excluded from analysis). *See* **Table S3**.

### The Sets of Proteins Targeted by PAV-951 and PAV-805 are Associated with the Hallmarks of Cancer

The proteins identified in PAV-951 and PAV-805 resin eluates by MSMS were searched in a cancer hallmarks database.^40^ The hallmarks database identifies gene sets which are associated with the ten classic hallmarks of cancer— replicative immortality, reprogramming energy metabolism, resisting cell death, tumor promoting inflammation, tissue invasion and metastasis, sustained angiogenesis, evading immune destruction, evading growth suppressors, genome instability, and sustaining proliferating signaling.^40^ Significant enrichment was found for the hallmarks of reprogramming energy metabolism and resisting cell death in the PAV-951 target (*see* **Figure 5S** and **Table S4** modified from Menyhart et. al.). Significant enrichment was found for the hallmark of reprogramming energy metabolism in the PAV-805 targets (*see* **Figures 5T** and Table S4, modified from Menyhart et. al.). Despite the overlap between the sets of proteins detected in the PAV-805 resin eluates from DMSO and PAV-805 treated cells, when the set of proteins detected by MSMS in the PAV-805 treated resin eluates were assessed in the cancer hallmarks enrichment analysis, no significant enrichment for any cancer hallmark was identified. *See* **Figures 5U** and Table S4, modified from Menyhart et. al.).

To examine whether our compounds have an effect on metabolic reprogramming in living cells rather than cell-extracts, MCF7 breast cancer cells were treated with DMSO, PAV-951, or PAV-805. After 20 hours of treatment, the concentration of glucose in the cell culture media was measured. A decrease in the cells’ glucose consumption was observed following PAV-951 and PAV-805 treatment. *See* **Figure 6**. The results provide evidence in support of the hypothesis that key features of metabolic reprogramming are disrupted with assembly modulator compound treatment.

**Figure 6.**
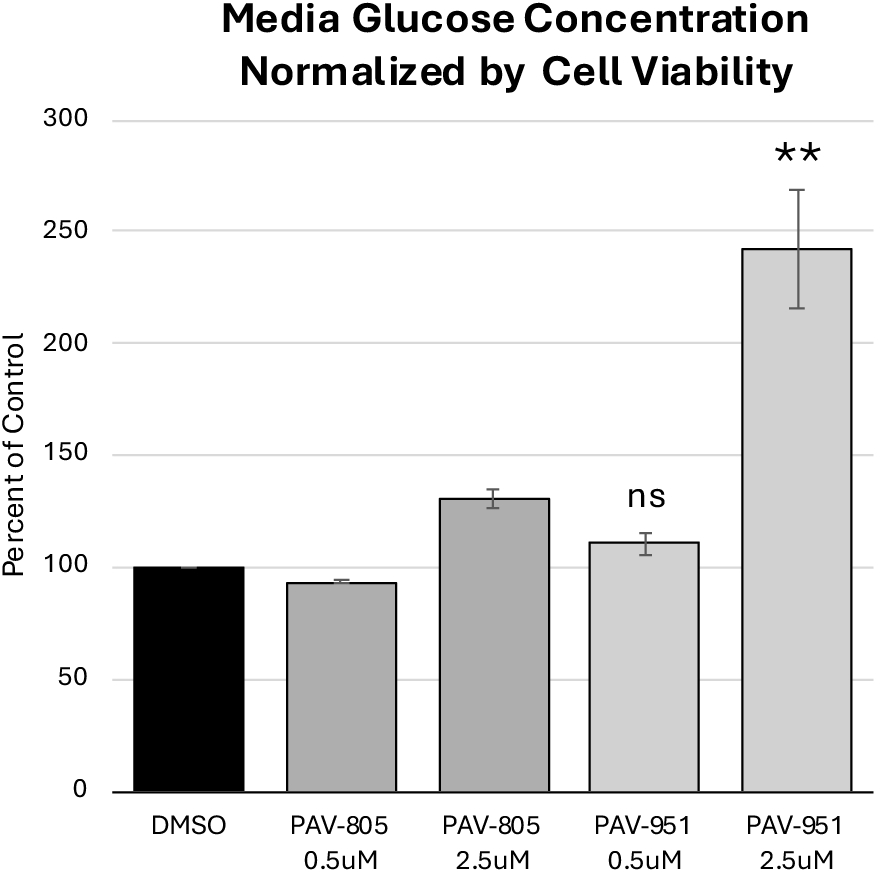
Treatment with PAV-805 and PAV-951 Decreases Glucose Utilization in Cancer Cells. MCF7 breast cancer cells were cultured in 5% fetal bovine serum media supplemented with 1.1mM glucose and treated with DMSO, 0.5uM, or 2.5uM of PAV-805 or PAV-951. After 20 hours, cell viability was measured by AlamarBlue^TM^ and the concentration of glucose in the media was measured by an Abcam glucose assay kit. Graphs shows percent increase of glucose in the cell culture media, relative to DMSO, normalized by cell viability. Error bars indicate deviation between samples. Statistical analysis was performed on the PAV-951 treated samples (as there were three replicates) using a t-test and the decreased glucose consumption observed between the DMSO and 2.5uM PAV-951 treated cells was found to be significant (p-value = 0.006).

### PAV-805 Binds Directly to PDI and indirectly to KAP1 In a Multiprotein Complex Target in Multiple Cancer Cell Extracts

A photocrosslinker analog of PAV-805, modified to contain a diazirine and a biotin group at the same position where resin had been attached, was synthesized to provide insight on which protein component was directly bound to the compound and which components were indirectly bound as part of a multi-protein complex held together through non-covalent interactions. *See* **Figure S6** for the chemical structure of the PAV-805 photocrosslinker analog.

The PAV-805-treated A549, HT29, and U251 extracts were incubated with the photocrosslinker analog, exposed to UV light, and precipitated with streptavidin under native and denatured conditions, with the denatured samples revealing the direct drug-binding protein(s) and the native samples revealing all proteins contained in the drug-binding complex. Western blot analysis identified PDI in both the native and denatured precipitations, and KAP1 in the native but not denatured precipitations for triplicate-repeated samples from all three cell types. *See* **Figure 7**.

**Figure 7.**
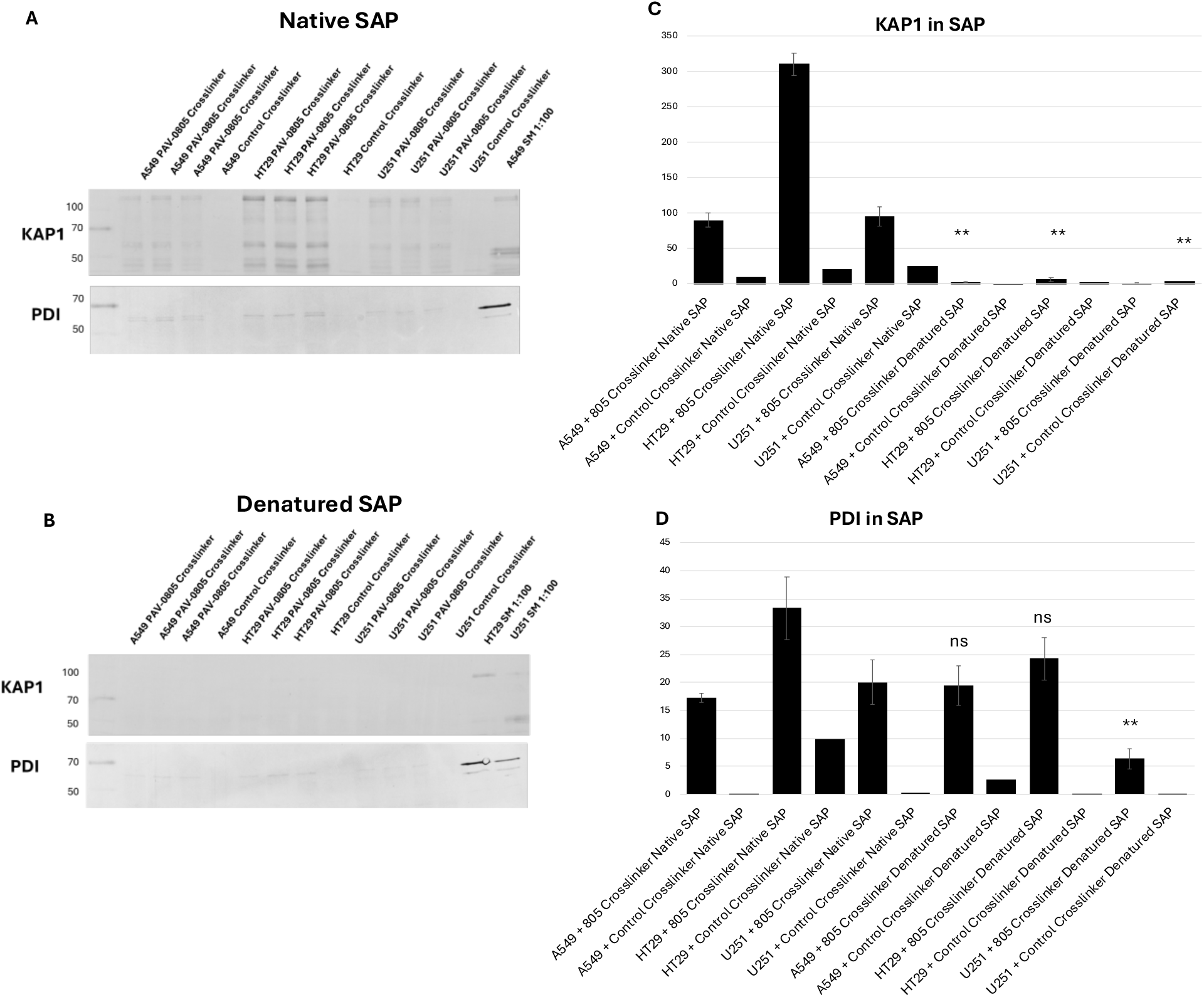
PAV-805 Directly Targets PDI in A549, HT29, and U251 Cell Extracts. A549, HT29, and U251 cells were treated with 1uM PAV-0805, and cell lysate was prepared. 70ul of 2 mg/ml PAV-805 treated cell extracts was incubated with 70ul of 2uM of a control photocrosslinker or a photocrosslinker analog of PAV-805 and then exposed to UV light for 10 minutes. The crosslinked samples were divided into two aliquots. One aliquot was kept native and one was denatured with addition of SDS and DTT, and boiled. Buffer containing excess Triton-X-100 was added to both aliquots, to adjust the denatured samples back to non-denaturing conditions. Strepatavidin beads were used to precipitate the biotinylated material. **Figures 5A** and **B** show western blots for KAP1 and PDI in the crosslinked streptavidin precipitations (SAP). **Figures 5C** and **D** shows quantitation of the protein bands, where triplicates have been averaged and error bars denote standard deviation.

### Molecular Modeling of the Compound Binding Site on PDI

In order to better understand the binding interactions of PAV-805 with PDI we performed *in silico* docking experiments with both PAV-951 and PAV-0805. The X-ray crystal structure of PDI is available at Protein Data Bank solved at 2.6 A (PDB: 3F8U).(44,84). Several possible binding pockets were predicted by the software utilized for docking experiments. Although both small molecules share a highly similar scaffold and the overall 3D shape, docking experiments suggest that they bind at the different sites and different orientations in the identified binding pockets. In silico experiments show that PAV-805 interacts with chain B, while PAV-951 engages more with Chain C of the heterodimer. *See* **Figure 8A**.

**Figure 8.**
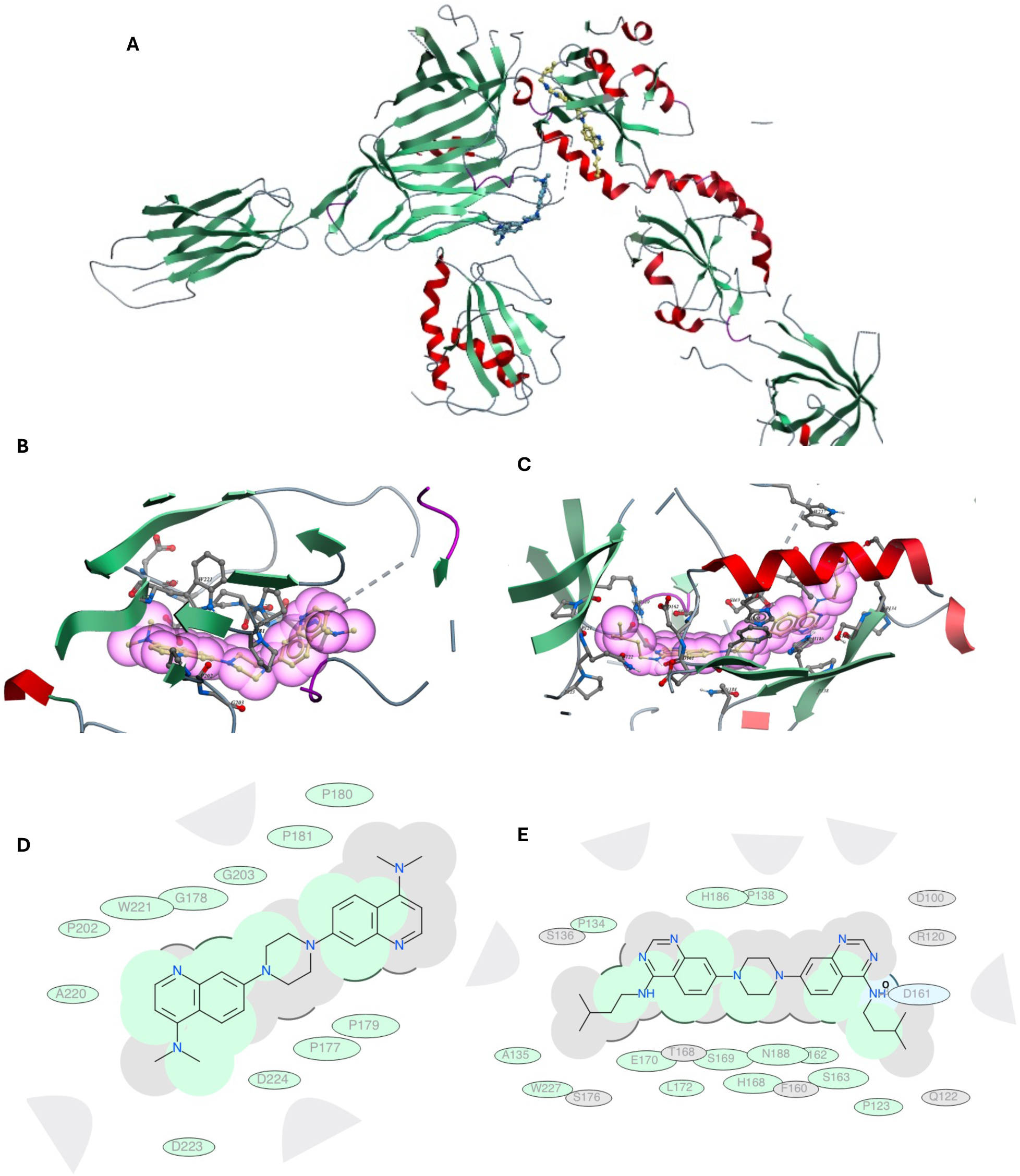
*In silico* Modeling of binding interactions between PAV-805, PAV-951, and PDI. **Figure 8A** shows binding poses of PAV-0805 (shown in blue) interacting with chain B, and PAV-951 (shown in yellow) binding to Chain C of the heterodimer. **Figures 8B** and **8C** show 3D representations of PAV-805 and PAV-951, respectively, within the binding site of the complex. Main amino acids in the proximity (below 5 A) of the ligands are shown. **Figures 8D** and **8E** show 2D representations of the binding pose of PAV-805 and PAV-951, respectively. Green shading represents hydrophobic region, gray parabolas represent accessible surface for large areas, blue represents hydrogen bonding, broken thick line around ligand shape indicates accessible surface, and the size of the residue ellipse represents the distance from the surrounding amino acid residues. **Table S5** shows the bond length differences and mf scores for the interactions of PDI with PAV-951 and PAV-805.

Analyzing the predicted PAV-805 binding interactions, we noticed several noncovalent interactions between quinolinyl moiety of PAV-805 with the residues A220, D223, D224, P202, G178, including π-π stacking with W221.The last two listed amino acid residues, G178 and W221 are in close proximity to the ligand. *See* **Figure 8** and **Table S5.**

Analyzing the predicted binding pose of PAV-951, we observed more interactions than with PAV-805, but of lower affinity. There is a hydrogen bond present between the 4-amino group (hydrogen on the nitrogen atom acts as a hydrogen bond donor) at the quinazolinyl ring and D161 (the oxygen atom is a hydrogen bond acceptor). The alkyl chain is interacting with Q122, S163, P123 and D162 via Van der Waals forces. The quinazoline ring is establishing hydrophobic interactions with D100, D162 and R120. One part of the piperazine ring is tightly surrounded with P138, S169 and N188, while the other side is opened towards the solvent which is shown by the absence of the thick line. The second quinazolinyl ring is even more engaged in hydrophobic interactions than the first ring, and is close to several amino acid residues, including T168, L172, H186 and P134. There are no π-π stacking with both quinazolinyl rings. *See* **Figure 8** and **Table S5.**

In contrast to the binding site identified by *in silico* docking for PAV-805 above, the active site of PDI has been identified at C57-G58-H59-C60 for the a domain and C406-G407-H408-C409 for the a’ domain (65, 69).

## Discussion

### SAR Exploration of Antiviral Assembly Modulating Series Leads to PAV-805, an Analog that Appears to Have Enhanced Target-Selectivity

The primary challenge for advancing oncology drugs is how to develop a compound that kills cancer cells and spares healthy cells. Protein heterogeneity– whether due to covalent post-translational modifications, intrinsically unfolded domains, or alternate pathways of protein biogenesis– is a dimension of gene expression that could hold the key to solving this challenge.^43–46^

Protein heterogeneity encompasses subsets of a particular protein that, for any of those reasons, are unique components of specific multi-protein complexes involved in specific functions. “Moonlighting” functions of proteins that are not readily identifiable through their sequence alone and reflect the functional side of the above (covalent or non-covalent) structural differences.^35,47–49^ Implicitly, from the concepts of protein heterogeneity (structural) and moonlighting (functional) it follows that only a small subset of the total amount of a protein in question that is found in a cell will be carrying out cancer-promoting functions. However, these subsets are what drives tumorigenesis and cancer development. A truly cancer-selective therapy would target this miniscule subset of the protein(s) that have departed from the maintenance of homeostasis. This therapy would be fundamentally non-toxic to a cell that is following its programming for homeostasis, as the molecular basis for collateral damage to healthy cells would be avoided *<u>because the target subset of the protein is not present in a healthy cell</u>*.

Viruses evolved the ability to achieve selectivity within the jungle of post-translational heterogeneity through natural selection in struggle with their hosts. Our phenotypic screen, carried out in a CFPSA system, started from where the viruses left off and identified compounds which modulate the assembly pathways that viruses evolved to exploit. A recurring theme we have observed with multiple assembly modulating compounds identified through our CFPSA screen for different viral families is that they appear to be able to selectively target particular subsets of proteins.^8,10,11,50,51^ Our compound PAV-104 appears to *<u>selectively</u>* target a subset of the protein 14-3-3 which is present in coronavirus-infected cells, but is not present in uninfected cells.^10^ Our compound PAV-866 appears to *<u>selectively</u>* target a subset of the protein ABCE1 which is found in fractions 2, 3, and 4 on a glycerol step gradient, despite ample ABCE1 protein being present across the gradient.^8^ Our compound PAV-617 appears to *<u>selectively</u>* target an oxidized form of MIF.^11^

Our findings must be taken in context that the functional impact of a protein is reflected in the “company that it keeps.” These particular subsets of proteins were identified by viruses as they evolved to exploit host machinery to catalyze assembly of their viral capsids.^5,8^ These subsets of proteins are likely strongly associated with disease states and likely carry out anti-homeostatic functions. The ability to to drug a protein only when found as part of a particular “aberrant” assembly complex could allow for only drugging a specific disease-implicated function of a multi-functional protein. Our fundamental hypothesis in approaching the work described in this paper, is that if there is an overlap between the subsets of proteins that cause dysregulation to homeostasis in a way that may both be exploited by viruses and cancer cells, then there is an opportunity to identify and advance an assembly-modulating cancer thereapeutic.

Our anti-viral assembly modulating compound PAV-951 showed broad but variable activity against the NCI-60 tumor cell panel and modest activity in the xenograft mouse models for A549 lung cancer and HT29 colon cancer.^1^ SAR exploration demonstrated that small changes to the chemical structure of PAV-951 could either eliminate or expand upon the anti-cancer activity (*See* **Figure 1)**. Multiple analogs were identified with no anti-cancer activity whatsoever. Multiple analogs were identified which were 100% lethal for every single tumor line examined, at doses non-toxic to healthy cells (*See* **Figure 1)**. Active analogs could be compared side-by-side, to assess potency relative to one another (*See* **Figure 2**). The emergence of a robust SAR indicated that the efficacy observed was structure-based and not non-specific.

Dose-response studies against tumor cell lines show that the compounds’ mechanism is both inhibitory on tumor cell proliferation and is lethal to the cancer cells, with what appears to be a window between the concentrations that are inhibitory to cancer cell growth (GI50) and the concentrations that are lethal to cancer cells (LC50). Dose-response studies in healthy PBMCs (which do not proliferate in cell culture), show that analogs from the PAV-951 series which show broad leathality to cancer cells in the NCI-60 screen are not lethal to these healthy cells even at high doses.

Due to limited resources, we chose to focus on compound safety in animals, rather than cell culture, on the rationale that animal safety is more comprehensive and characterizing the therapeutic window between safety and efficacy *in vivo* is more relevant for guiding decisions on translatability in humans. PAV-951 could be safely administered to mice at a dose of 2.5 mg/kg.^1^ Improved animal safety relative to the parent compound, was observed for some but not all of the analogs which displayed improved pan-cancer activity in the NCI screen, indicating that toxicity to healthy animals is not an inherent feature of this particular anti-cancer mechanism *See* **Figures 1** and **2**.

PAV-805 is the safest compound identified from the PAV-951 series thus far. Mice display no adverse affects following twice-daily administration of PAV-805 at 10 mg/kg for two weeks. Meanwhile, once-daily administration of 10 mg/kg PAV-805 (i.e. half the safe limit for multi-day dosing) was efficacious in inhibiting tumor growth and metastasis in early and late stage presentations of metastatic breast cancer in the 4T1 mouse allograft model.

The enhanced safety and breadth of the anti-cancer activity observed with of PAV-805 compared to the parent compound, PAV-951, is consistent with our general model that assembly modulating compounds access a specific disease-associated protein target within the milieu of post-translational heterogeneity and that SAR can be advanced to increased selectivity for that specific subset of protein.^8,10,11,50^ Increased selectivity to a subset of protein that is only present in dysregulated cells, is one plausible explanation for our data showing that the efficacy against cancer cells increased in tandem with improvements to mouse safety.

### 4T1 Allograft Achieved Proof-of-Concept for Anti-Cancer Efficacy of Advanced Compounds Against Triple Negative Metastatic Breast Cancer

Nontoxic doses of PAV-620, PAV-621, and PAV-805 significantly inhibited tumor growth and reduced metastasis in the 4T1 mouse allograft model for triple negative metastatic breast cancer. The anti-cancer activity of the compounds in the mouse model was comparable to Paclitaxel (Taxol), which is an FDA-approved first-line treatment for metastatic breast cancer in humans.^52^ The activity of PAV-621, as superior to Paclitaxel in inhibiting tumor growth, was replicated across two 4T1 allograft studies. Compared side-by-side on the metric of tumor growth inhibition, PAV-621 was more efficacious than PAV-620, which was more efficacous to PAV-805. However, all three compounds (and the positive control Paclitaxel), were essentially equal on the metric of reduction of metastasis. By acting upon metastasis in addition to reducing the primary tumor, these compounds appeared to be superior to other compounds probed against the 4T1 model in the literature which only had an effect on inhibiting tumor growth^53^.

One variable examined in the two allograft studies was the time that compound treatment began. In the first study, PAV-621 was equally potent when administered beginning at the time ot tumor implantation versus at the time when the tumor had grown to 50mm^3^. In the second study, PAV-805 was equally potent beginning when the tumor was at a size of 50mm^3^ and 500mm^3^. In both studies, the late treatment groups showed an immediate effect on inhibiting tumor growth, and by the end of the study had essentially “caught up” to the early treatment groups on both metrics of tumor size and number of metastases. We interpret these findings to suggest that the mechanism is equally active on *both* early and late-stage cancers *in vivo*. Together with the compound’s novel mechanism of action, these findings suggest PAV-805 or another analog in the series could be developed into a viable treatment for patients with late-stage cancers that have either not responded to or have developed resistance against currently available chemotherapies.

The dose of compound administered in the PAV-620 treatment group was lowered from 5 mg/kg to 1.5 mg/kg after one week of treatment due to safety concerns, as the multi-day safety study conducted prior to the allograft indicated that daily 5 mg/kg dosing is only well-tolerated for one week. However, despite having only received one week of treatment at a dose which would be predicted to yield plasma concentrations sufficient to reach EC50, on day 21 both the size of the primary tumor and the number of metastases were significantly reduced, relative to control. We interpret these findings to suggest that a short period of treatment can elicit significant and long-lasting anti-cancer effects, so long as sufficient exposure is achieved during the treatment window. This may reduce the risk of long-term toxic side effects in a clinical context, where patients would only need to take high doses of the compound for short period of time.

### PAV-620 and PAV-805 Demonstrate a Novel Mechanism in Primary Carcinomas

Following the allograft success, we used primary breast cancer cells to probe the mechanism of compounds in the PAV-951 series. We were interested in observing the compounds’ effect under conditions that trigger anoikis, a particular form of programmed cell death that occurs when cells detach from their extracellular matrix.^38,39^ Cancer cells can only metastasize to distant sites once they collect the mutations needed to bypass anoikis.^38,39^ Our interest in anoikis was suggested by early data that PAV-951 is significantly more cytotoxic to cells that have been released from substrate binding by trypsin, a condition that is known trigger anoikis.^1,34,39^ The literature indicates that oncogenic viruses drive anoikis resistance in infected cells and therefore we thought it plausible that if our compounds normalize changes at the host-viral interface, their mechanism may restore anoikis to metastasizing cells.^54,55^ We hypothesized that elevated programmed cell death by this mechanism could contribute to the efficacy of the PAV-951 series against metastasis.

Increased cytotoxicity to primary cells from breast carcinomas was observed following a short period of PAV-620 and PAV-805 treatment of primary carcinoma cells seeded under conditions that should induce anoikis. *See* **Figure 4**. Cells treated for the same amount of time with Paclitaxel or Etoposide did not show cell death above the baseline amount induced by anoikis. *See* **Figure 4**. We interpret the results to suggest that carcinoma cells have a particular sensitivity to PAV-805 and PAV-620 under conditions that trigger anoikis. Due to limited bandwidth, we have not yet parsed out whether the compounds’ observed cytotoxicity is due to a shared or distinct mechanism from its observed inhibition on cancer cell proliferation.

2-dimensional UMAP analysis of images of the primary breast carcinoma cells showed PAV-620 and PAV-805 treated cells clustering together in a distinct region from cells treated with the apoptosis inducing agent Staurosporine.^56^ *See* **Figure 3**. Unexpectedly, cells treated with Paclitaxel and Etoposide clustered together with the negative control DMSO. *See* **Figure 3**. We speculate that since clustering by UMAP frequently corresponds to shared mechanisms, the phenotypic effects of anoikis were dominant over other potential features due to the drugs targeting cell division (Paclitaxel) and DNA damage (Etoposide).^57,58^. From the UMAP, we concluded that PAV-620 and PAV-805 are cytotoxic to carcinomas cells in which anoikis has been triggered likely via the same mechanism-of-action as one another, and a different mechanism from those of Staurosporine, Paclitaxel, or Etoposide.

### Characterizing Target of PAV-805 Compared to Target of PAV-951

Results from photocrosslinking and eDRAC experiments showed that, like its parent compound PAV-951, PAV-805 targets a dynamic multi-protein complex.^1^ Photocrosslinking in lung, colon, and brain cancer samples revealed that PAV-805 binds directly to PDI. *See* **Figure 7**. Other proteins, including KAP1, are targeted by PAV-805 indirectly as components of the multi-protein complex. *See* **Figure 7**. When the eDRAC eluates containing the target complexes of PAV-951 and PAV-805 were compared and they were found to contain overlapping, but distinct sets of proteins. *See* **Figure 5**. One notable difference is that the target of PAV-951 appears to include component proteins of the proteasome, while the target of PAV-805 does not. By MSMS, all three triplicate-repeated PAV-951 resin eluate samples contained 11 different proteasome component proteins while the PAV-805 resin eluate samples did not contain any proteasome component proteins.^1^ Western blots of DRAC run in parallel with the PAV-951 and PAV-805 resins confirmed this finding. *See* **Figure 5**.

We speculate that the difference in protein composition detected in the PAV-805 target may indicate increased selectivity of the compound for particular subsets of proteins that are found in particular multi-protein complexes. Inclusion of the proteasome proteins in the PAV-951 target and not the PAV-805 target may indicate that the mechanism of PAV-951 includes a second function that the mechanism of PAV-805 does not, or it could indicate that PAV-951 has a greater propensity for off-target effects due to targeting multi-protein complexes which are not necessary for anti-cancer activity, or both. Evidence for the first hypothesis includes the hallmarks enrichment analysis indicating that PAV-951 targets sets of proteins implicated in two cancer hallmarks while PAV-805 only targets sets of proteins implicated in one cancer hallmark. Evidence for the second hypothesis includes the improvements to animal safety observed with PAV-805 as increased selectivity towards a particular target complex may correspond to decreased toxicity, as the potential for “collateral damage” of off-target effects decreases in tandem.

The selectivity of a given analog for a particular PDI-containing multi-protein complex is highly relevant within our framework for understanding assembly-modulating compounds. PDI proteins are multi-functional and involved in numerous protein-protein interactions^59,60^. PDI proteins are found in different cellular compartments including the endoplasmic reticulum (ER) and the nucleus, and are secreted from the ER.^60,61^ The literature describes PDI carrying out distinct catalytic activities that include facilitating the isomerization of disulfide bonds, the oxidation of thiols, and oxidative protein folding.^60,62,63^ PDI is described in the literature as undergoing a number of post-translational modifications.^62,64,65^ Some of these modifications are linked to particular diseases including, for example, that S-nitrosylation of PDI is associated with neurodegeneration and S-glutathionylation of PDI is associated with cancer.^64–66^ PDI has also been observed to associate with the HIV viral protein Env in the context of viral entry and assembly.^63,67–69^ Upregulation of PDI is observed in a variety of different cancers, and in metastasis.^70^ Inhibition of PDI enhances ferroptosis, an iron-dependent form of programmed cell death. ^71^ Paglia et. al., showed that inhibition of PDIA3, the particular PDI isoform identified in the PAV-805 target, in glioblastoma cells led to arrest at the G1/S checkpoint and showed a redistribution of PDIA3 which included visible formation of protein aggregates.^72^ To the extent that protein aggregates may also be viewed as protein assemblies, the associations between PDIA3 inhibition, formation of assembly complexes, and phenotypic observations showing an arrest of cancer cell growth is likely relevant to the mechanism of compounds in the PAV-951 series. More work is needed to investigate the specific role of PDI and PDI-containing protein assembilies in the compounds’ anti-cancer mechanism.

### Characterizing the PAV-805 Target in Multiple Cell Types and in Response to Compound Treatment

We used western blot analysis to compare the amounts of four critical proteins of the cancer interactome, ALDH1A1, KAP1, PDI, and YWHAZ, which were found in the PAV-805 DRAC eluate generated from starting material prepared from lung, colon, and brain cancer. While KAP1, PDI, and YWHAZ were detected in all eluates, ALDH was detected by western blot in the PAV-805 resin eluate from lung and colon cancer cell lysate, but not in the resin eluate from brain cancer cell lysate *See* **Figure 5**. This indicates that the target complex may have a different composition from one cancer cell to another. This is consistent with an upstream, allosteric mechanism of action where compound treatment restores an aberrant assembly machine to its normal function/composition regardless of which proteins have gone awry as part of the cancer’s cellular reprogramming. This may help to explain the broad activity against diverse cancers as the compounds may normalize aberrant biochemical pathways that are driven selectively by each of a range of cancer-causing triggers by analogy to the evidence that these compounds are effective on all members of a given viral family and in some cases on multiple viral families.

When DMSO versus PAV-805 treated lung, colon, and brain cancer cell extracts were compared on the PAV-805 resin, a 100kDa KAP1 band was detected by western blot in the PAV-0805-treated target. *See* **Figure 5**. We thought the enrichment of this particular form of KAP1 in the target with PAV-805 treatment was notable given the described roles of KAP1 in the literature. KAP1 is an allosteric modulator involved in a variety of protein-protein interactions an array of functions including transcriptional activation of HIV, T-cell development, DNA damage repair, as a transcriptional co-repressor for many genes, and as a ligase for post translational modifications such as ubiquitination and SUMOlyation.^73,74^ The diversity of functions that KAP1 displays appears to be, at least in part, through its assembly into different multi-protein complexes that carry out different objectives.^31,37,75–77^ That treatment with PAV-805 triggers that recruitment of a *<u>particular</u>* 100 kDa form of KAP1 into the PAV-805 target, may be a critical step in the compound’s anti-cancer mechanism.

MSMS analysis of eDRAC eluates showed that when A549 cells were treated with PAV-805, the proteins HSPA5, PDIA3, ALDH3A1, ALDH1A1, AHCY, and CKM, which had been detected in the DMSO-treated eluates, remain in the PAV-805 target. *See* **Table S3**. Meanwhile, the proteins AMY2A and AHNAK, not previously detected in DMSO-treated eluates, appear in the PAV-805 target. *See* **Table S3**. The proteins detected in both the DMSO and PAV-805 treated eluates have been associated with cancer in the literature or are used as diagnostic factors^70,70,78–81^. Some of the proteins, such as AHCY and CKM, are described as carrying out both pro-cancer or cancer-suppressive functions, depending on the context of what protein-protein interactions that they are involved in.^80,81^ Of the proteins that appear with PAV-805 treatment, there is scant literature connecting AMY2A with cancer, though one study noted it may play a role as a tumor suppressor gene.^82^ Meanwhile, AHNAK is broadly described as playing a role in cancer, including roles suppressing cancer development and metastasis.^83^

Our working model of the PAV-805 mechanism, based on results from eDRAC experiments, is that compound treatment recruits particular tumor-suppressive proteins into a multi-protein complex and that drives multi-functional protein components to selectively engage in pro-homeostatic functions through their inclusion in the target complex. This mirrors what we have observed for other, structurally-unrelated, assembly modulating compounds, including in other therapeutic areas, that were identified through the original CFPSA screen.^1,10,51^. While we have not yet been able to fully characterize the significance of particular protein components within the target complex, we look forward to continuing to build on these discoveries because we hypothesize that the information content revealed by the protein-protein interactions involving *<u>cellular subsets</u>* of proteins that are targeted by assembly modulating compounds cannot be readily identified by more conventional approaches, including machine-learning. A challenge for these future studies is that the small subset of PDI involved is a barrier to selective genetic manipulation (knockdown, overexpression, or resistant mutants) in order to establish the causal relationship between complex remodeling and anti-cancer activity.

### Characterizing the Binding Site Through *in silico* Modeling

*In silico* docking studies showed that PAV-951 and PAV-805 bind PDI at slightly different locations with different affinities. *See* **Figure 8**. PAV-805 bound with higher affinity despite fewer interactions. PAV-805 is only predicted to interact with PDI exclusively through hydrophobic interactions, while PAV-951 was predicted to interact with PDI through both hydrophobic interactions and hydrogen bonding. The differences in the predicted binding site for PAV-951 and PAV-805 is consistent with other observations on the differences between the two molecules including that the set of proteins targeted by PAV-805 is smaller, the observed activity of PAV-805 is selective for only one hallmark of cancer, and the safety of PAV-805 is much improved, relative to PAV-951.

The predicted binding site is different from the active site of PDI as described in the literature. ^70,84,85^ This is consistent with our hypothesis that *<u>assembly modulating compounds have allosteric targets</u>*. An allosteric compound is one that can regulate a protein’s action from a location distinct from its active site. In theory, an allosteric drug would act in a more physiological manner than most conventional pharmacological compounds as it allows for upstream regulation of a protein’s activity. Compounds acting allosterically may display more selective effects on the behavior of their target protein(s) rather than broad inhibition of the protein’s catalytic activity, which is what most target-based methods of drug discovery are specifically designed to do. The advancement from PAV-951 to PAV-805 through SAR exploration provides evidence that it is possible to optimize an allosteric compound to be more selective in its target, which may correspond to downstream improvements including reduced toxicity and greater potency.

### Connecting Protein Assembly to the Hallmarks of Cancer

The MSMS results revealing enrichment in specific cancer hallmark-associated gene products for PAV-951 and PAV-805 resin eluates provides a framework for understanding how cancer-associated biochemical pathways could be impacted by assembly modulation. *See* **Figure 5**. The enrichment analysis, which is an association based on the literature and not empirically demonstrated through experimentation, showed that the set of proteins found to be targeted by PAV-951 were associated with the cancer hallmarks of resisting programmed cell death and reprogramming energy metabolism, while the set of proteins found to be targeted by PAV-805 were only associated with the cancer hallmark of reprogramming energy metabolism.

Resisting programmed cell death is a fundamental hallmark of cancer.^32,86^ Restoring a crucial control point that allow inappropriately proliferating cells to be driven to programmed cell death provides a novel, broad, and potent anti-cancer mechanism. The set of proteins in the PAV-951 eluate that were enriched for resisting programmed cell death included the proteosome proteins which were not targeted by PAV-805. *See* **Figure 5**. The results suggest that early assembly modulating compounds may have multiple targets, but can be driven towards selectivity for one versus others. The change in both predicted affinity and position within the binding site suggests that those alterations could account for the loss by PAV-805 of association with one of the two cancer hallmarks seen in PAV-951.

Reprogramming energy metabolism is the second cancer hallmark significantly enriched in the PAV-951 drug resin eluates, that remains enriched in the PAV-805 target. The Warburg Effect describes a metabolic reprogramming where cancer cells shift towards use of glycolysis over oxidative phosphorylation for energy generation, even in the presence of oxygen.^41^ Activity against abnormal metabolism pathways provides an alternate explanation for the potent activity of the compounds in metastasis models, as metabolic reprogramming is a critical component of the EMT.^41^ The Warburg Effect is implicated in viral infection as well, and it allows both the viruses and cancer to bypass homeostasis ^16,25–30^.

When DRAC was conducted using cancer cells that had been treated with PAV-805 (rather than DMSO) as starting material, fewer proteins were observed in the target by MSMS. While there was significant overlap in the eluate composition (all but two of the proteins found in triplicate-repeated samples of the PAV-805 treated eluates where also found in the triplicates of the DMSO-treated eluates), the hallmarks analysis on the compound-treated eluates no longer showed that the set of targeted proteins were enriched for any cancer hallmarks. Within the framework that we have developed to understand the mechanism of assembly modulator compounds, we interpret these results to suggest that treatment with PAV-805 changed the composition of an aberrant assembly complex into that of a normal assembly complex. While the normal complex contains many of the same proteins as the aberrant one, *<u>its function within the cell has changed and it no longer serves to drive cancer progression</u>*.

### Preliminary Data Show Changes to Metabolism in Cell Culture Following Compound Treatment

The cancer hallmarks enrichment analysis associating the compounds’ drug target to metabolic programming in cancer cells prompted us to examine glucose consumption in cancer cells treated with PAV-951 and PAV-805. Treatment of breast cancer cells with PAV-951 and PAV-805 increases the concentration of glucose in the cell culture media, indicating a *<u>decrease</u>*in glucose consumption in response to compound treatment. *See* **Figure 6**. These results suggest that association between the proteins targeted by PAV-951 and PAV-805 and the hallmark of reprogramming metabolism is functional, that is the compounds’ mechanism acts to counter pro-cancer metabolic programming, such as the Warburg Effect, in some way.

A puzzling observation is that the effect was greater for PAV-951 than for PAV-805. As these studies were carried out at a single time point, it is possible that the time course of the effect is different for the different compounds. Profiling the effects of more time points and more compounds within the SAR set on glucose consumption in cancer cells would provide valuable insight. The studies also must be expanded on to examine the effect that compounds have on other metabolic products, such as lactic acid. Further studies will be needed to determine which metabolic pathways are up-regulated and down-regulated in response to compound treatment and to parse out the difference between the changes in glucose consumption induced by treatment with PAV-951 compared to PAV-805. Metabolic flux studies could be conducted to further define whether the compounds’ effect on glucose consumption represents glycolytic suppression, mitochondrial restoration, or broader metabolic remodeling.

## Future studies

These studies show that assembly modulating compounds from the PAV-951 chemical series are broadly efficacious in cell and mouse models for cancer. Further work, including off-target liability profiling and GLP PK/toxicology studies, is needed to optimize this series as a potential human cancer therapeutic. Further work is also required and provide more insight to the multi-protein complex target and mechanism of action. Although these studies identify a PDI-containing assembly complex associated with cancer metabolism as the target of PAV-805, the temporal sequence connecting complex remodeling, metabolic changes, and induction of cell death remains to be fully established.

Nevertheless, we are encouraged that our compounds show the potential for development into novel, first-in-class cancer therapeutics and as biochemical tools for examining otherwise inaccessible protein-protein interactions. We originally identified PAV-951 due to its modulation of HIV capsid assembly in a CFPSA system ^1^ If viruses evolved to exploit post-translational heterogeneity when reprogramming host machinery to assemble capsids and trigger disease, we hypothesize that assembly modulating compounds identified through our screen can access the same dimension of post-translational heterogeneity and catalyze protein assembly as a means for reversing disease. The interactions that associate PAV-951 and its analogs to PDI and PDI to the other proteins in the target, could reveal novel information on the molecular basis for homeostasis.

## Methods

### National Cancer Institute 60 Cell Line Screen

In the NCI-60 screen, up to 60 cancer cell lines (CCRF-CEM, HL-60(TB), K-562, MOLT-4, RPMI-8226, SR, A549/ATCC, EKVX, HOP-62, HOP-92, NCI-H226, NCI-H23, NCI-H322M, NCI-H460, NCI-H522, COLO 205, HCC-2998, HCT-116, HCT-15, HT29, KM12, SW-620, SF-268, SF-295, SF-539, SNB-19, SNB-75, U251, LOX IMVI, MALME-3M, M14, MDA-MB-435, SK-MEL-2, SK-MEL-28, SK-MEL-5, UACC-257, UACC-62, IGROV1, OVCAR-3, OVCAR-4, OVCAR-5, OVCAR-8, NCI/ADR-RES, SK-OV-3, 786-0, A498, ACHN, CAKI-1, RXF 393, SN12C, TK-10, UO-31, PC-3, DU-145, MCF7, MDA-MB-231/ATCC, HS 578T, BT-549, T-47D, MDA-MB-468) are grown.

Two methods have been used to assess anti-tumor cytotoxicity in the NCI-60 screen.

In the Classic Screen (method used through September 2023), cells were grown for 24 hours then treated with vehicle or compound, or fixed in situ with TCA, to represent a measurement of the cell population for each cell line at the time of drug addition. The treated cells were grown for an additional 48 hours before being fixed in situ with TCA. Fixed cells were then stained with Sulforhodamine B. Absorbance was read to determine cell viability of compound-treated cells relative to both the time at which treatment began and to untreated cells at the end of the study. The NCI-60 screen was initially conducted at a single dose of 2.5uM, but was subsequently repeated with a dose titration.

The human tumor cell lines of the cancer screening panel are grown in RPMI 1640 medium containing 5% fetal bovine serum and 2 mM L-glutamine. For a typical screening experiment, cells are inoculated into 96 well microtiter plates in 100 μL at plating densities ranging from 5,000 to 40,000 cells/well depending on the doubling time of individual cell lines. After cell inoculation, the microtiter plates are incubated at 37° C, 5 % CO2, 95 % air and 100 % relative humidity for 24 h prior to addition of experimental drugs.

After 24 h, two plates of each cell line are fixed in situ with TCA, to represent a measurement of the cell population for each cell line at the time of drug addition (Tz). Experimental drugs are solubilized in dimethyl sulfoxide at 400-fold the desired final maximum test concentration and stored frozen prior to use. At the time of drug addition, an aliquot of frozen concentrate is thawed and diluted to twice the desired final maximum test concentration with complete medium containing 50 μg/ml gentamicin Following drug addition, the plates are incubated for an additional 48 h at 37°C, 5 % CO2, 95 % air, and 100 % relative humidity. For adherent cells, the assay is terminated by the addition of cold TCA. Cells are fixed in situ by the gentle addition of 50 μl of cold 50 % (w/v) TCA (final concentration, 10 % TCA) and incubated for 60 minutes at 4°C. The supernatant is discarded, and the plates are washed five times with tap water and air dried. Sulforhodamine B (SRB) solution (100 μl) at 0.4 % (w/v) in 1 % acetic acid is added to each well, and plates are incubated for 10 minutes at room temperature. After staining, unbound dye is removed by washing five times with 1 % acetic acid and the plates are air dried. Bound stain is subsequently solubilized with 10 mM trizma base, and the absorbance is read on an automated plate reader at a wavelength of 515 nm. For suspension cells, the methodology is the same except that the assay is terminated by fixing settled cells at the bottom of the wells by gently adding 50 μl of 80 % TCA (final concentration, 16 % TCA).

In the HTS384 Screen (method used after September 2023), the sixty human tumor cell lines of the cancer screening panel are grown in RPMI 1640 medium containing 5% fetal bovine serum and 2 mM L-glutamine. To initiate a screen, 40 μl of cells are inoculated into the wells of white 384-well microtiter plates at plating densities ranging from 250 to 2,500 cells/well, depending on the doubling time of individual cell lines. After inoculation, the microtiter plates are incubated at 37°C with 5% CO^2^ and 95% relative humidity for 24 h before the addition of controls and test agents. Prior to a screen, the test agents and controls are solubilized in dimethyl sulfoxide (DMSO) at 400-fold the final test concentration and stored frozen in polypropylene 384-well microtiter plates.

Twenty-four hours after cell line inoculation, acoustic dispensing is used to transfer 100 nl of DMSO (0.25% (v/v), final) into the wells of microtiter plates, each containing a single cell line. Subsequently, 40 μl of CellTiter-Glo are dispensed into the wells, according to the manufacturer’s protocol. Luminescence is measured to assess cell viability at the time of drug addition (time zero, Tz). Acoustic dispensing is also used to transfer 100 nl of controls and test agents (400-fold dilution, final) into duplicate microtiter plates for each cell line to achieve technical replicates. Controls in each microtiter plate include vehicle, 100% cytotoxicity (1 μM Staurosporin NSC755774 and 3 μM Gemcitabine NSC613327 [n = 8]), and five concentrations of doxorubicin (NSC123127, 25 μM [n = 2], 2.5 μM [n = 1], 250 nM [n = 2], 25 nM [n = 1], 2.5 nM [n = 2]. Following the delivery of controls and test agents, the microtiter plates are incubated for 72 h at 37°C with 5 % CO^2^ and 95% relative humidity. After 72 h of exposure, 40 μl of CellTiter-Glo are dispensed into the wells of the microtiter plates and luminescence is measured, according to the manufacturer’s protocol, to assess cell viability.

For both methods, using the seven absorbance measurements [time zero, (Tz), control growth, (C), and test growth in the presence of drug at the five concentration levels (Ti)], the percentage growth is calculated at each of the drug concentrations levels. Percentage growth is calculated as:

[(Ti-Tz)/(C-Tz)] x 100 for concentrations for which Ti>/=Tz

[(Ti-Tz)/Tz] x 100 for concentrations for which Ti<Tz.

Three dose response parameters are calculated for each experimental agent. Growth inhibition of 50 % (GI50) is calculated from [(Ti-Tz)/(C-Tz)] x 100 = 50, which is the drug concentration resulting in a 50% reduction in the net protein increase (as measured by SRB staining) in control cells during the drug incubation. The drug concentration resulting in total growth inhibition (TGI) is calculated from Ti = Tz. The LC50 (concentration of drug resulting in a 50% reduction in the measured protein at the end of the drug treatment as compared to that at the beginning) indicating a net loss of cells following treatment is calculated from [(Ti-Tz)/Tz] x 100 = -50.

PAV-951, PAV-0383, PAV-0384 were assessed in the Classic Screen. All others were assessed by the HTS384 screen.

### Cytotoxicity Assessment on A549 and HT29

A549 and HT29 cells were seeded at a density of 5,000 cells/well and incubated at 37oC for 24 hours. Cells were then treated with either DMSO or compound, at concentrations of 20uM, 5uM, 1.25uM, or 0.325uM. The cells were incubated at 37oC for 72 hours then cell viability was measured using an Titer-Glo assay.

### Cytotoxicity Assessment on PBMCs and H9 Lymphoma

Healthy human PBMCs and H9 cells were seeded at a density of 5,000 cells/well and treated with either DMSO or compound, at concentrations of 2.5uM, 5uM, 10uM, and 20uM. The cells were incubated at 37oC for 72 hours then cell viability was measured using an AlamarBlue^TM^ assay.

### Mouse Safety

For the single-dose MTD study, female Balb/c mice aged 8-10 weeks were randomly divided into treatment groups with three animals per group. Animals in each treatment group were weighed and received one IP injection of 0.1-0.15mL containing either vehicle (10% DMSO, 45% propylene glycol, 45% sterile water), or varying doses compound. Animals were observed from for 24 hours for clinical signs of toxicity. MTD was determined to be the dose at which no signs of toxicity were observed by any parameters.

For multi-day safety studies, compounds were administered to three male and three female BALB/c mice by IP dosing once or twice daily and animals were monitored daily for adverse effects.

### Mouse Allograft

A suspension of 0.1×10^6^ cells were implanted into the mammary fat pad of normal balb/c mice. Size of the tumor was measured using a digital Vernier caliper. Tumor volume was calculated as the [Length (L) × Width (W) ^2^] /2, where length (L) is the largest diameter of the tumor and width (W) is the smallest diameter. Tumor growth inhibition was calculated based on the following formula:

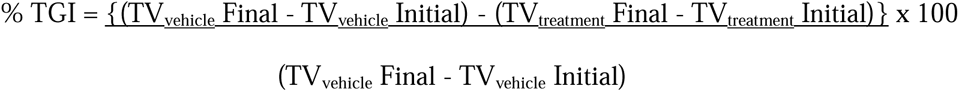

In the first allograft study, animals were treated by IP injection, beginning on the same day as tumor establishment, with either vehicle (N=6), Paclitaxel (N=6), or PAV-621 (N=6). An additional late treatment group (N=6) received vehicle until the tumor volume in the vehicle group reached 50-60mm^3^, In the second allograft study, animals were treated by IP injection, beginning on the day when the tumor had grown to a volume of 50mm^3^, with either vehicle (N=6), 2 mg/kg Paclitaxel (N=6), 5 mg/kg then (after 8 days) 1.5 mg/kg PAV-620, 2.5 mg/kg PAV-621, or 10mg/kg PAV-0805 (N=6), beginning when the tumor volume reached 50mm^3^. An additional late treatment group (N=3) received vehicle until the tumor volume reached 500mm^3^, and were subsequently treated with 10mg/kg PAV-0805 daily. Throughout both studies, animals were continually weighed and monitored for adverse clinical signs. In both studies, animals were sacrificed once the tumor in the vehicle-treated group reached a volume of 2000mm^3^. Lung tissues were isolated and visually observed for metastatic cancer cell colonies. In the first allograft study, the following hematology studies were conducted upon completion of the allograft: Total Leucocyte Count, Total Erythrocyte Count, Percent Haematocrit, Percent Hemoglobyn, MCV, MCH, MCHC, Platet Count, Urea, ALT, AST, CRE, Percent Neutrophils, Percent Lymphocytes, Percent Monocytes, Percent Eosinophils, and Percent Basophils.

### Assessment in Primary Breast Carcinoma

BB5 and BB7 cells were seeded at 2000 cells/well into PhenoPlate 384-well microplates (Cat # 6057302, Revvity) pre-treated with PolyHEMA coating. Briefly, 20ul of 20mg/mL PolyHEMA was added to each well, then allowed to air dry with the lid open in a biosafety cabinet to ensure sterility. After drying, the same amount was added to the wells a second time. Once dried, the plate was used for anoikis assays. Drug compounds were added along with seeded BB5 and BB7 cells. Live cell stains ChromaLIVE non-toxic dye (Saguaro Biosciences) and Annexin-V labelled with Alexa-647 (made in house) were added to the cells upon seeding. Each concentration had a duplicate as a technical replicate. DMSO conditions had 6 technical replicates. At 24 hours post-treatment, Hoechst 33342 at 1µg/mL was added and the cells were imaged on a Revvity Opera Phenix screening confocal microscope. A total of 16 fields-of-views were imaged for each well. Harmony image analysis software was used to post-process the images. A total of 293 features related to intensity, morphology, and texture features were extracted. A random forest model was generated in python from the package sklearn, using sampled images of cells (from both BB5 and BB7 cell types) from the DMSO condition as ‘DMSO’, and 2.5µM Staurosporine as ‘Dead’. Normalized area under the curve was calculated as the total AUC after the baseline death percentage (percent of dead cells at 0µM) was removed from all data points.

### Glucose Assay

Glucose concentrations in cell culture supernatants were determined using the L-lactate assay kit (Abcam, #65331) and the glucose assay kit (Abcam, #65333), respectively, in accordance with the manufacturer’s protocols.

MCF7 cells (2 × 10 per well) were seeded in 96-well plates. After 24 h, the medium was replaced with phenol red–free DMEM supplemented with 5% dialyzed fetal bovine serum (FBS) and 1.1 mM glucose. Cells were treated with the indicated compounds and incubated for 20 h. Supernatants were subsequently collected and directly analyzed for lactate and glucose content.

Cell viability was evaluated using a resazurin (Alamar Blue) assay. Briefly, 10 µL of a 100 µg/mL resazurin solution was added to each well, followed by incubation for 2 h at 37 °C. Absorbance was measured at 570 nm with a reference wavelength of 620 nm.

### Drug Resin Affinity Chromatography

A549, HT29, or U251 cells were grown in minimum essential media (UCSF) with 10% FBS and 1% Penstrep for 48 hours. In some conditions cells were treated with PAV-0805 for 20 hours. Cells were scraped into cold phosphate buffered saline (PBS) (10mM sodium phosphate, 150 mM sodium chloride pH 7.4), then spun at 1,000 rpm for 10 minutes until pelleted. The PBS was decanted and the pellet resuspended in a low salt buffer (10mM HEPES pH 7.6, 10mM NaCl, 1mM MgAc with 0.35% Tritonx100) then centrifuged at 10,000 rpm for 10 minutes at 4o C. The post-mitochondrial supernatant was removed and adjusted to a concentration of approximately 5 mg/ml unless otherwise stated and equilibrated in a physiologic column buffer (50 mM Hepes ph 7.6, 100 mM KAc, 6 mM MgAc, 1 mM EDTA, 4mM TGA). The extract was supplemented with an energy cocktail (to a final concentration of 1mM rATP, 1mM rGTP, 1mM rCTP, 1mM rUTP, creatine phosphate and 5 ug/mL creatine kinase). 30 ul of extract was then incubated for one hour at 22oC on 30 ul of affi-gel resin coupled to PAV-951, PAV-0805, or a 4% agarose matrix (control). The input material was collected and the resin was then washed with 3 ml column bufffer. The resins were eluted overnight at 22oC in 100ul column buffer containing 100uM PAV-951 or PAV-0805 with the energy cocktail. Eluate samples were sent for MSMS analysis.

### Chemical Photocrosslinking

Cell extract was prepared as above then adjusted to a protein concentration of approximately 2 mg/ml in column buffer (50 mM Hepes pH 7.6, 100 mM KAc, 6 mM MgAc, 1 mM EDTA, 4mM TGA). 70ul of 2uM stock of photocrosslinker analogs of PAV-805 chemically modified to contain biotin and a diazirine group or a control containing the biotin and diazarine but not assembly modulator compound prepared in column buffer were added to 70ul of extract, incubated for one hour at on ice, then exposed to ultraviolet light for 10 minutes on ice. After crosslinking, samples were divided in two 60 ul aliquots and one set was denatured by adding 6ul of 10% SDS, 0.75 ul DTT, and boiling for 10 minutes. Both native and denatured aliquots were then diluted in 1200 ul column buffer containing 0.1% triton. 5 ul of magnetic streptavidin beads were added to all samples and mixed for one hour at room temperature to capture all biotinylated proteins and co-associated proteins. Samples were placed on a magnetic rack to hold the beads in placed and washed three times with 1mL of column buffer containing 0.1% triton. After washing, beads were resuspended in 40 ul of gel loading buffer containing SDS and analyzed by western blot for PDI and KAP1.

### Silver Stain

SDS/PAGE gels were incubated overnight in a fixative (50% methanol, 10% acetic acid, 40% water), then for an hour in 50% methanol (done as two washes), and an hour in water (done as two washes). The gels were sensitized in 0.02% sodium thiosulfate for one minute then washed twice for 30 seconds with water. The gels were incubated for 30 minutes in cold 0.1% silver nitrate with 0.02% formaldehyde then washed twice for 30 seconds. The gels were developed in 3% sodium carbonate with 0.02% formaldehyde. The developed gels showing the pattern of protein bands was scanned and the image was analyzed.

### Western blotting

SDS/PAGE gels were transferred in Towbin buffer (25mM Tris, 192mM glycine, 20% w/v methanol) to polyvinylidene fluoride membrane, blocked in 1% bovine serum albumin (BSA) in PBS, incubated overnight at 4 degrees Celsius in a 1:1,000 dilution of 100ug/mL affinity-purified primary IGG to PDI, HIV Gag, KAP1, MDH, YWHAZ in 1% BSA in PBS containing 0.1% Tween-20 (PBST). Membranes were then washed twice in PBST and incubated for two hours at room temperature in a 1:5000 dilution of secondary anti-rabbit or anti-mouse antibody coupled to alkaline phosphatase in PBST. Membranes were washed two more times in PBST then incubated in a developer solution prepared from 100 uL of 7.5 mg/mL 5-bromo-4-chloro-3-indolyl phosphate dissolved in 60% dimethyl formamide (DMF) in water and 100ul of 15 mg/ml nitro blue tetrazolium dissolved in 70% DMF in water, adjusted to 50mL with 0.1 Tris (pH 9.5) and 0.1 mM magnesium chloride. Membranes were scanned and the integrated density of protein band was measured on ImageJ. Averages and the standard deviation between repeated experiments were calculated and plotted on Microsoft Excel.

### MSMS Analysis

Samples were processed by SDS PAGE using a 10% Bis-ttris NuPAGE gel with the MES buffer system. The mobility region was excised and washed with 25 mM ammonium bicarbonate followed by 15mM acetonitrile. Samples were reduced with 10 mM dithoithreitol and 60 degrees Celsius followed by alkylation with 5o mM iodacetamide at room temperature. Samples were then digested with trypsin (Promega) overnight (18 hours) at 37 oC then quenched with formic acid and desalted using an Empore SD plate. Half of each digested sample was analyzed by LC-MS/MS with a Waters NanoAcquity HPLC system interfaced to a ThermoFisher Q Exactive. Peptides were loaded on a trapping column and eluted over a 75uM analytical column at 350 nL/min packed with Luna C18 resin (Phenomenex). The mass spectrometer was operated in a data dependent mode, with the Oribtrap operating at 60,000 FWHM and 15,000 FWHM for MS and MS/MS respectively. The fifteen most abundant ions were selected for MS/MS.

Data was searched using a local copy of Mascot (Matrix Science) with the following parameters: Enzyme: Trypzin/P; Database: SwissProt Human (concatenated forward and reverse plus common contaminants); Fixed modification: Carbamidomethyl (C) Variable modifications: Oxidation (M), Acetyl (N-term), Pyro-Glu (N-term Q), Deamidation (N/Q) Mass values: Monoisotopic; Peptide Mass Tolerance: 10 ppm; Fragment Mass Tolerance: 0.02 Da; Max Missed Cleavages: 2. The data was analyzed by label free quantitation (LFQ) and spectral count methods. LFQ intensity values of each condition were measured in triplicate and compared against each other to generate log2 fold change values for each combination of conditions. Spectral counts were filtered for a 1% protein/peptide false discovery rate requiring 2 unique peptides per protein. The data set was further adjusted by subtraction of spectral counts for specific proteins observed in the control resin, and subtraction of proteins that were not found in all triplicate-repeat samples. The adjusted dataset was searched for enrichment of cancer hallmarks in the cancerhallmarks.com dataset as described in https://doi.org/10.1016/j.jpha.2024.101065.

### Molecular Modeling

Docking studies were performed using ICM Pro (Molsoft LLC). A human protein disulfide isomerase (PDI) model was prepared from the crystal structure (PDB ID: 3F8U). The 3F8U file was imported into ICM, converted to an ICM object, and energy-minimized using default program settings. Putative binding pockets were then identified using the ICM pocket-finding algorithm with standard search parameters.

PAV-805 and PAV-951 were subsequently docked into the selected pocket(s) according to the ICM docking workflow. Docking was conducted with a thoroughness setting of 50 (effort per ligand), generating three conformations per ligand. Docked poses were evaluated using the resulting Mean Force scores (mf scores) - the potential of mf score evaluates the strength of interactions based on statistical preferences of atom-atom contacts. The lower mf score, i.e. more negative score represents a better, more energetically stable predicted binding pose.

## Authors Contributions

AFL: conceptualization, investigation, methodology, data curation, project administration, supervision, writing-original draft, SM: data curation, investigation, project administration, MM: investigation, data curation, methodology DS: conceptualization, JL: investigation, methodology, MDP: supervision, DD: supervision, DS: conceptualization, methodology, BBL: investigation, methodology, AMS: investigation, methodology, DWA: supervision, CK: supervision, SP: investigation, methodology, VRL: conceptualization, investigation, methodology, supervision, writing-review and editing.

## Acknowledgements

We thank Amanda Macieik, Shao Feng Yu, Anatoliy Kitaygorodskyy, Kumar Paulvannan, Umesh Appaiah, Dmitry Temnikov, Louisa Ng, and Andrew Braham for being part of this work. We thank the National Cancer Institute for running the NCI-60 screening data shown. Funding for these studies was provided by Prosetta Biosciences.

## Figure Captions

**Figure S1. NCI-60 Dose Titrations.** Dose titrations of PAV-620 and PAV-805 were assessed against the panel of cancers in the NCI-60 screen. **Figure S1 A-I** show dose-response curves of the tumor cell lines in response to PAV-620 treatment, grouped by cancer type. **Figure S1 1J-R** show dose-response curves of the tumor cell lines in response to PAV-805 treatment, grouped by cancer type. **Supplemental Figure 1S** shows calculated IC50 values for each treatment condition and tumor cell line.

**Table S1. Multi-day Safety of PAV-621, PAV-620, and PAV-805.** PAV-621, PAV-620 and PAV-805 were administered to male and female BALB/c mice by IP dosing once or twice daily for two weeks and animals were monitored daily for adverse effects. Observations of normal are indicated with the letter “N” and observations of lethargy are indicated with the letter “L”. PAV-621 was administered daily at 2.5 mg/kg. No adverse effects were observed with PAV-621 administration. Initially, PAV-620 was administered twice daily at 5 mg/kg. Animals appeared normal for the first two days. On day 3, all animals appeared lethargic. On day 4, the dosing schedule was decreased to a single IP dose of 5 mg/kg daily and the study continued. Animals appeared normal from day 4 through day 11. On day 12, all animals appeared lethargic. The animals recovered on day 13 and dosing resumed. The animals appeared normal for the remainder of the study. PAV-805 was administered twice daily at 10 mg/kg. No adverse effects were observed with PAV-805 administration.

**Table S2. Mouse PK of PAV-621, PAV-620 PAV-805. 5 mg/kg of PAV-621,** PAV-620 and PAV-805 were administered to CD1 mice by IP injection. Plasma was collected at 8 time points over the subsequent 24 hours and the concentration of compound in the plasma was measured at each time point in ng/mL.

**Figure S3. Body Weight of Animals in First 4T1 Allograft Study.** 4T1 mouse breast cancer tumors were transplanted onto mice and treated with vehicle, Paclitaxel, or PAV-621. **Figure S3** shows the daily body weights of the mice during the study.

**Figure S4. Hematology of Animals in First 4T1 Allograft Study.** 4T1 mouse breast cancer tumors were transplanted onto mice and treated with vehicle, Paclitaxel, or PAV-621. **Figure S4** shows hematology parameters measured upon completion of the study. No treatment related variations were observed.

**Figure S5. Body Weight of Animals in Second 4T1 Allograft Study.** 4T1 mouse breast cancer tumors were transplanted onto mice and treated with vehicle, Paclitaxel, PAV-620, PAV-621, or PAV-805. **Figure S5** shows the daily body weights of the mice during the study.

**Figure S6. Structures of Resin and Photocrosslinker Analogs.** PAV-951 and PAV-805 were coupled to affi-gel resins for eDRAC experiments. A modified version of PAV-805 containing diazirine and biotin groups was synthesized for photocrosslinking experiments. **Figure S6** shows the resin attachment points and the chemical structure of the photocrosslinker compounds used in this paper.

**Figure S7.** Silver Stain and Western Blots from eDRAC Experiments. A549, HT29, and U251 cells were treated with DMSO or PAV-805 and lysate was prepared and used for eDRAC, repeated in triplicate on the PAV-951 and PAV-805 resins, as well as in single-point on a control resin. The eDRAC eluates were run on SDS PAGE and analyzed by western blot and MSMS. **Figure S7A** shows the silver stain protein pattern of the eluates from DMSO-treated A549 cell lysate run on the PAV-951, PAV-805, and control resins. **Figure S7B** shows western blots of the eluates from A549 cell lysate run on the PAV-951, PAV-805, and control resins. **Figure S7C** shows western blots of the eluates from DMSO and PAV-805 treated A549, HT29, and U251 cells run on the PAV-805 and control resins.

**Table S3. Proteins Identified in the PAV-805 Eluate by MSMS.** A549 cells were treated with DMSO or 1uM PAV-0805, and cell lysate was prepared and used for eDRAC, repeated in triplicate on PAV-805 resins, as well as in single-point on a control resin. The eluates were run on SDS PAGE and analyzed by MSMS. Table shows the number of spectral counts of given proteins for PAV-805 resin eluates in triplicate next to the matched sample eluted from the control resin in single point. Conditional formatting on a red-to-green scale has been applied to visualize the spectral counts of particular proteins relative to others in the dataset.

**Table S4. Enrichment for Cancer Hallmarks in the PAV-951 and PAV-805 Resin Eluate Proteins.** The sets of proteins in the PAV-951 and PAV-805 resin eluates were searched in a database for sets of genes associated with the hallmarks of cancer.^40^ For each cancer hallmark, chart shows the overlap between the list of proteins from the eluate and the database of hallmark-associated proteins, the p-value and odds ratio for association with the hallmark, and list of the hallmark-associated gene products detected in each set of proteins.

**Table S5. Predicted Contacts and Distances of PAV-951 and PAV-805 to PDI.** Molecular modeling experiments conducted *in silico* showed the predicted contacts and distances between PAV-951 and PAV-805 and residues within PDI.

