## Supplemental Figure 1 for "Structure-Activity-Relationship Advancement, Animal Validation, and Target Engagement of a Novel Assembly Modulator Small Molecule Chemical Series with Pan-Cancer Selective Cytotoxicity"

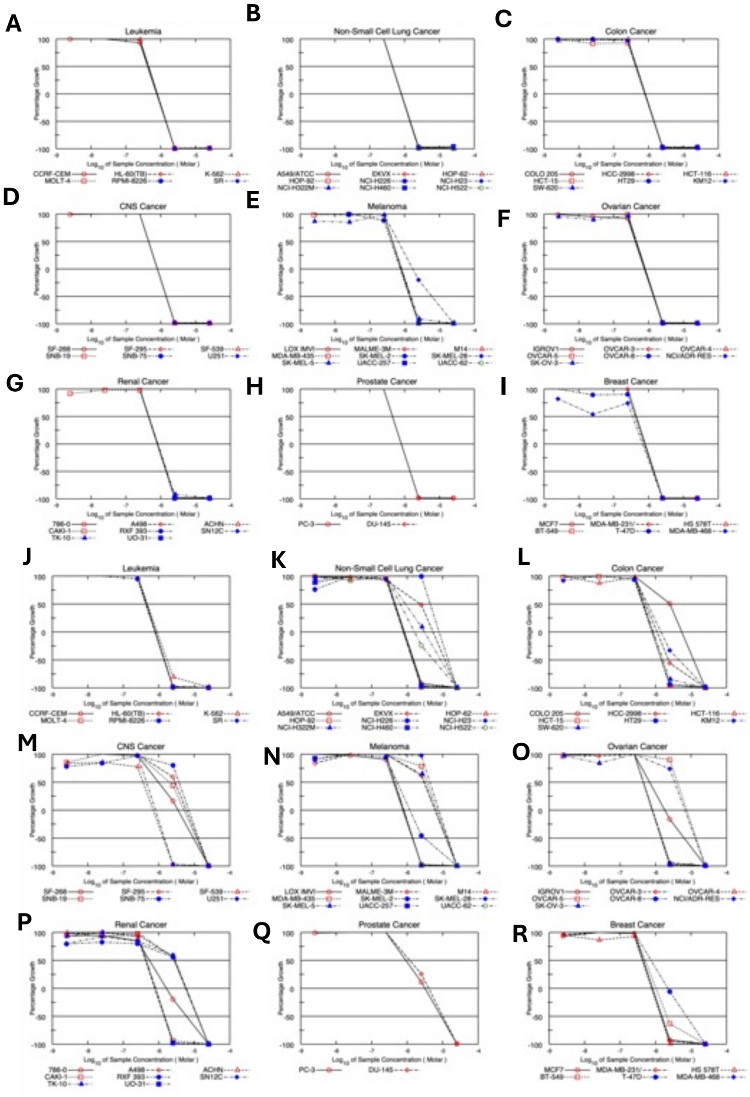


**S**

| Cancer Type | Cell Line | PAV-620 IC50 Log10 | PAV-805 IC50 Log10 |
|---|---|---|---|
| Leukemia | CCRF-CEM | -6.13 | -6.04 |
| | HL-60(TB) | -6.02 | -6 |
| | K-562 | -6.11 | -6.09 |
| | MOLT-4 | -6.06 | -6.04 |
| | RPMI-8226 | [redacted] | -6.12 |
| | SR | -6.07 | -6.08 |
| Lung Cancer | A549/ATCC | -6.06 | -6.04 |
| | EKVX | -6.05 | -5.15 |
| | HOP-62 | -6.05 | -6.09 |
| | HOP-92 | -6.05 | -6.11 |
| | NCI-H226 | -6.02 | -6.1 |
| | NCI-H23 | -6.08 | -6.12 |
| | NCI-H322M | -6.08 | -5.68 |
| | NCI-H460 | -6.03 | -6.1 |
| | NCI-H522 | -6.08 | -5.82 |
| Colon Cancer | COLO 205 | -6.07 | -5.42 |
| | HCC-2998 | [redacted] | -6 |
| | HCT-116 | -6.1 | -6.11 |
| | HCT-15 | -6.13 | -6.09 |
| | HT29 | -6.09 | -6.13 |
| | KM12 | -6.11 | -5.97 |
| | SW-620 | -6.06 | -6.06 |
| CNS Cancer | SF-268 | -6.08 | -5.59 |
| | SF-295 | -6.08 | -5.28 |
| | SF-539 | -6.02 | -6.2 |
| | SNB-19 | -6.07 | -5.39 |
| | SNB-75 | [redacted] | -5.14 |
| | U251 | -6.09 | -6.08 |
| Melanoma | LOX IMVI | [redacted] | -6.14 |
| | MALME-3M | -6.07 | -5.17 |
| | M14 | -6.08 | -6.08 |
| | MDA-MB-435 | -6.08 | -5.18 |
| | SK-MEL-2 | -6.14 | -5.94 |
| | SK-MEL-28 | -5.58 | -5.11 |
| | SK-MEL-5 | -6.09 | -5.26 |
| | UACC-257 | -6.07 | -6.06 |
| | UACC-62 | -6.07 | -6.04 |
| Ovarian Cancer | IGROV1 | -6.13 | -5.91 |
| | OVCAR-3 | -6.04 | -6.05 |
| | OVCAR-4 | -6.03 | -6.03 |
| | OVCAR-5 | -6.1 | -5.13 |
| | OVCAR-8 | -6.03 | -6.06 |
| | NCI/ADR-RES | -6.07 | -5.21 |
| | SK-OV-3 | -6.11 | -6.09 |
| Renal Cancer | 786-0 | -6.09 | -5.89 |
| | A498 | -6.05 | -6.13 |
| | ACHN | -6.07 | -5.34 |
| | CAKI-1 | -6.11 | -6.1 |
| | RXF 393 | -6.08 | -5.21 |
| | SN12C | -6.06 | -5.27 |
| | TK-10 | -6.07 | -5.28 |
| | UO-31 | -6.09 | -6.09 |
| Prostate Cancer | PC-3 | -6.08 | -5.8 |
| | DU-145 | -6.05 | -5.71 |
| Breast Cancer | MCF7 | -6.09 | -6.11 |
| | MDA-MB-231/ATCC | -6.1 | -6.06 |
| | HS 578T | -6.06 | -6.12 |
| | BT-549 | -6.08 | -5.97 |
| | T-47D | -6.12 | -5.65 |
| | MDA-MB-468 | -6.12 | [redacted] |
