## Supplementary figures and images for "Structure-Activity-Relationship Advancement, Animal Validation, and Target Engagement of a Novel Assembly Modulator Small Molecule Chemical Series with Pan-Cancer Selective Cytotoxicity"

### Supplemental Figure 2

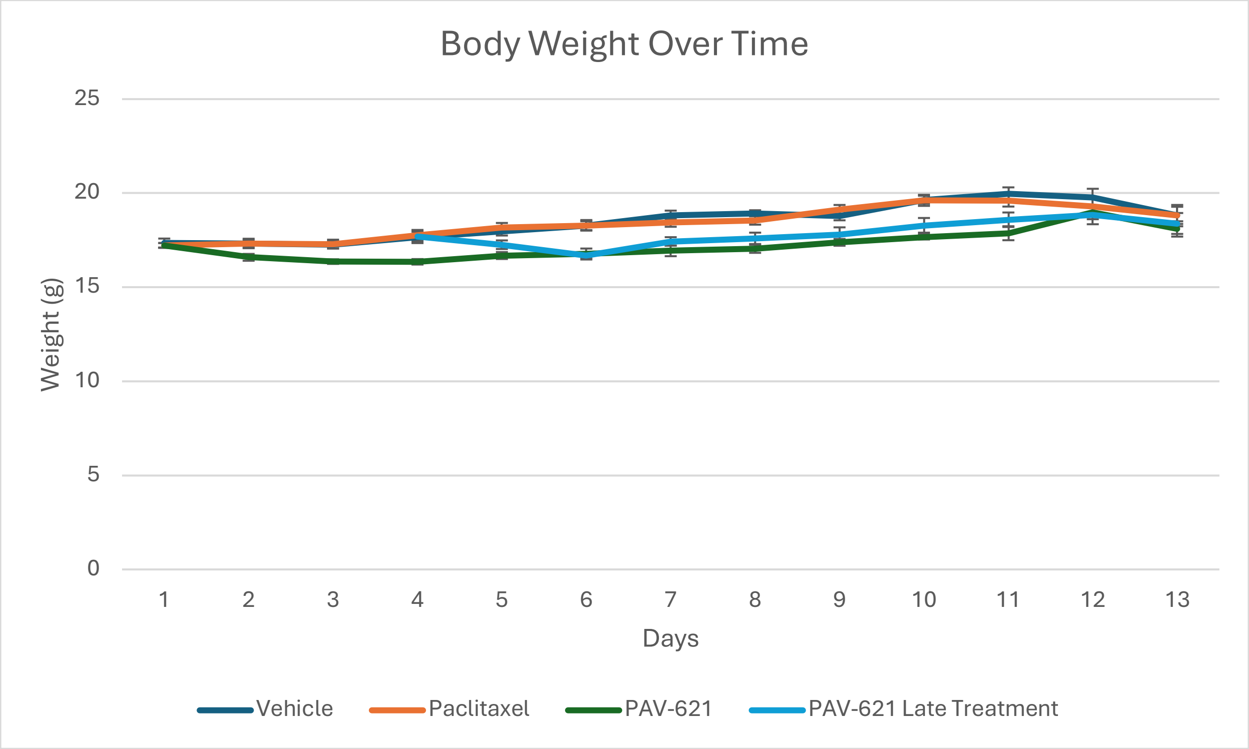

Body Weight Over Time
25
20
15
10
5
0
Weight (g)
1
2
3
4
5
6
7
8
9
10
11
12
13
Days
Vehicle
Paclitaxel
PAV-621
PAV-621 Late Treatment

### Supplemental Figure 4

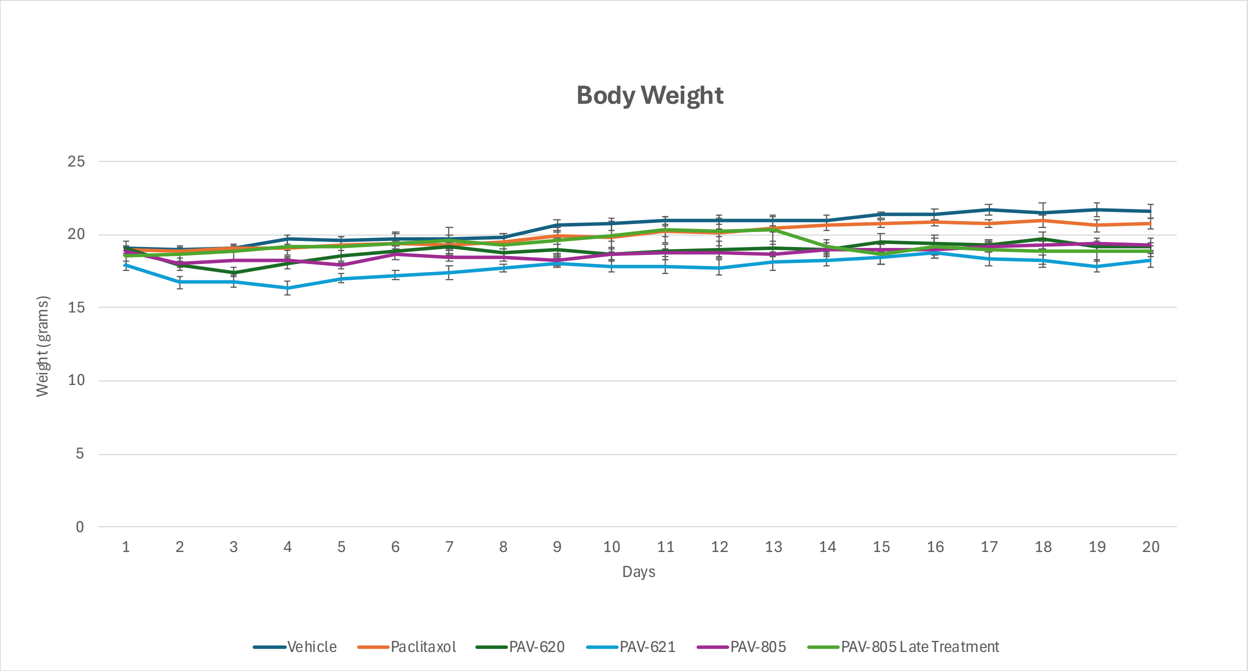

Body Weight
25
20
15
10
5
0
Weight (grams)
1
2
3
4
5
6
7
8
9
10
11
12
13
14
15
16
17
18
19
20
Days
Vehicle
Paclitaxol
PAV-620
PAV-621
PAV-805
PAV-805 Late Treatment

### Supplemental Figure 5

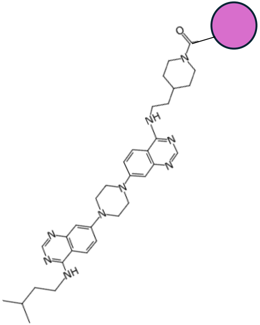

**PAV-951 Resin**

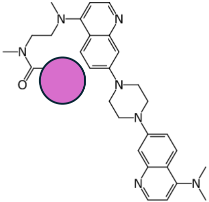

**PAV-805 Resin**

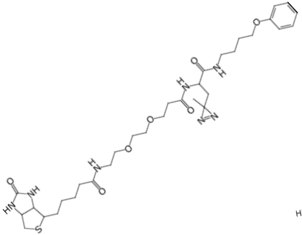

**Control Crosslinker**

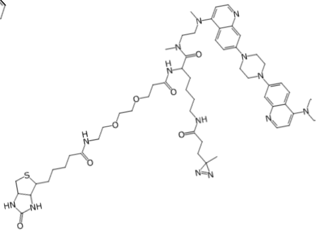

**PAV-805 Crosslinker**
