## Supplemental Figure 3 for "Structure-Activity-Relationship Advancement, Animal Validation, and Target Engagement of a Novel Assembly Modulator Small Molecule Chemical Series with Pan-Cancer Selective Cytotoxicity"

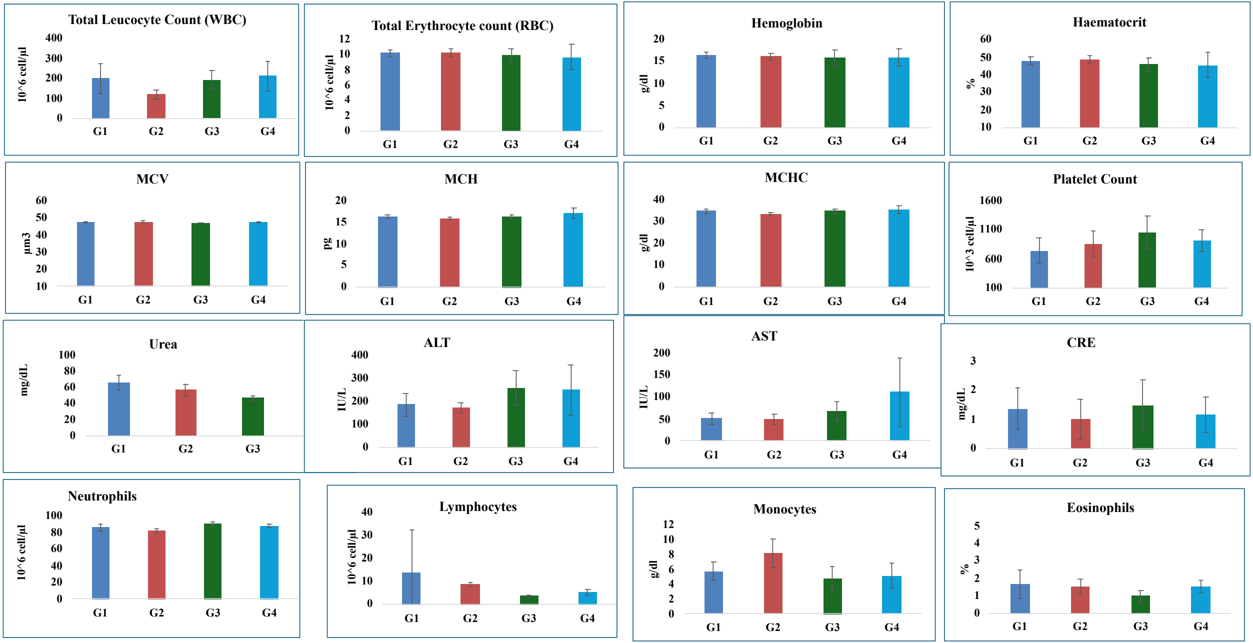
Total Leucocyte Count (WBC)
10^6 cell/µl
400 300 200 100 0
G1 G2 G3 G4
Total Erythrocyte count (RBC)
10^6 cell/µl
12 10 8 6 4 2 0
G1 G2 G3 G4
Hemoglobin
g/dl
20 15 10 5 0
G1 G2 G3 G4
Haematocrit
%
60 50 40 30 20 10
G1 G2 G3 G4
MCV
µm3
60 50 40 30 20 10
G1 G2 G3 G4
MCH
pg
20 15 10 5 0
G1 G2 G3 G4
MCHC
g/dl
40 30 20 10 0
G1 G2 G3 G4
Platelet Count
10^3 cell/µl
1600 1100 600 100
G1 G2 G3 G4
Urea
mg/dL
100 80 60 40 20 0
G1 G2 G3
ALT
IU/L
400 300 200 100 0
G1 G2 G3 G4
AST
IU/L
200 150 100 50 0
G1 G2 G3 G4
CRE
mg/dL
3 2 1 0
G1 G2 G3 G4
Neutrophils
10^6 cell/µl
100 80 60 40 20 0
G1 G2 G3 G4
Lymphocytes
10^6 cell/µl
40 30 20 10 0
G1 G2 G3 G4
Monocytes
g/dl
12 10 8 6 4 2 0
G1 G2 G3 G4
Eosinophils
%
5 4 3 2 1 0
G1 G2 G3 G4
