## Supplemental Figure 6 for "Structure-Activity-Relationship Advancement, Animal Validation, and Target Engagement of a Novel Assembly Modulator Small Molecule Chemical Series with Pan-Cancer Selective Cytotoxicity"

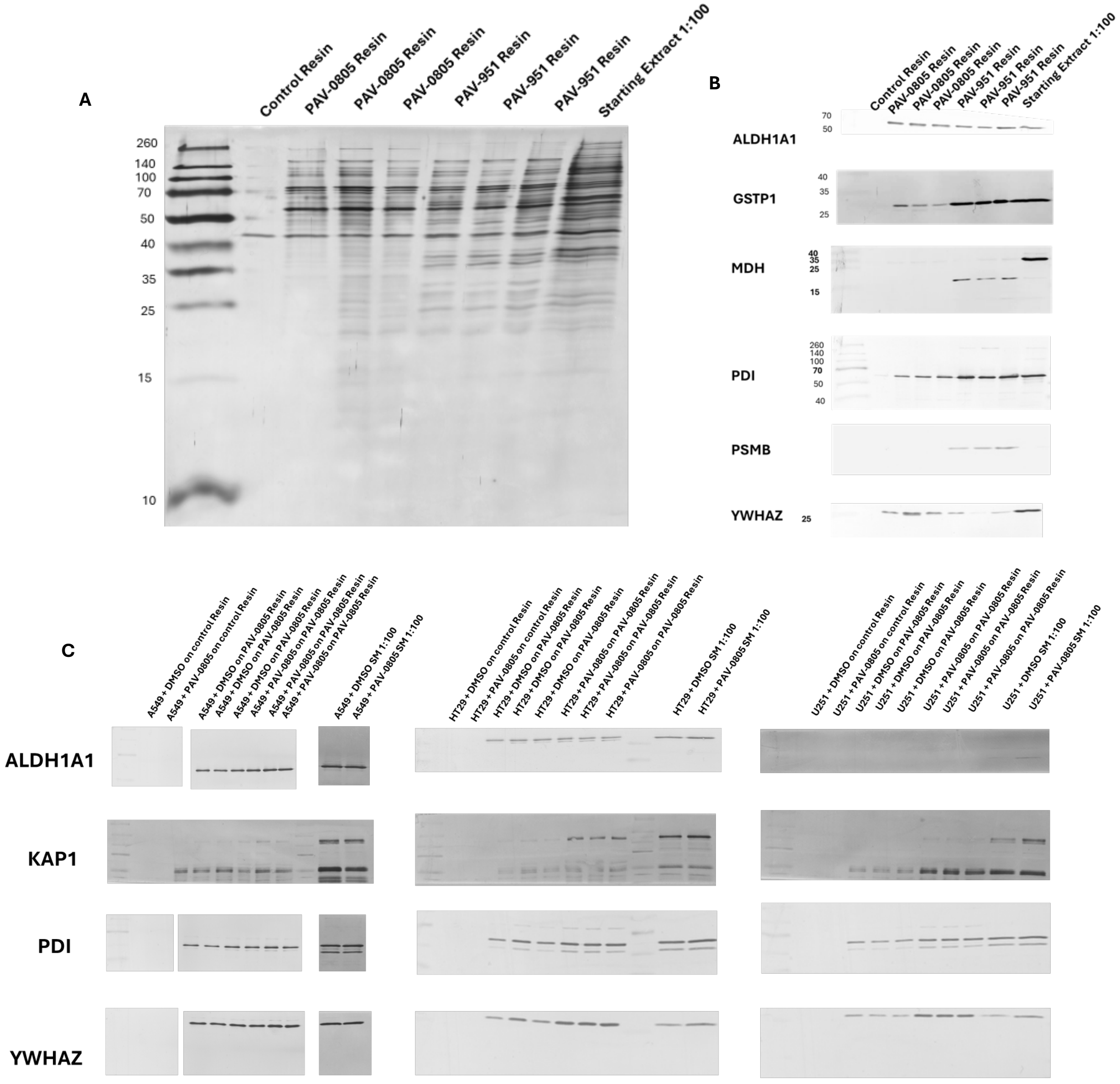
A
Control Resin
PAV-0805 Resin
PAV-0805 Resin
PAV-0805 Resin
PAV-951 Resin
PAV-951 Resin
PAV-951 Resin
Starting Extract 1:100
260
140
100
70
50
40
35
25
15
10
B
Control Resin
PAV-0805 Resin
PAV-0805 Resin
PAV-0805 Resin
PAV-951 Resin
PAV-951 Resin
PAV-951 Resin
Starting Extract 1:100
ALDH1A1
70
50
GSTP1
40
35
25
MDH
40
35
25
15
PDI
260
140
100
70
50
40
PSMB
YWHAZ
25
C
A549 + DMSO on control Resin
A549 + PAV-0805 on control Resin
A549 + DMSO on PAV-0805 Resin
A549 + DMSO on PAV-0805 Resin
A549 + DMSO on PAV-0805 Resin
A549 + PAV-0805 on PAV-0805 Resin
A549 + PAV-0805 on PAV-0805 Resin
A549 + PAV-0805 on PAV-0805 Resin
A549 + DMSO SM 1:100
A549 + PAV-0805 SM 1:100
HT29 + DMSO on control Resin
HT29 + PAV-0805 on control Resin
HT29 + DMSO on PAV-0805 Resin
HT29 + DMSO on PAV-0805 Resin
HT29 + DMSO on PAV-0805 Resin
HT29 + PAV-0805 on PAV-0805 Resin
HT29 + PAV-0805 on PAV-0805 Resin
HT29 + PAV-0805 on PAV-0805 Resin
HT29 + DMSO SM 1:100
HT29 + PAV-0805 SM 1:100
U251 + DMSO on control Resin
U251 + PAV-0805 on control Resin
U251 + DMSO on PAV-0805 Resin
U251 + DMSO on PAV-0805 Resin
U251 + DMSO on PAV-0805 Resin
U251 + PAV-0805 on PAV-0805 Resin
U251 + PAV-0805 on PAV-0805 Resin
U251 + PAV-0805 on PAV-0805 Resin
U251 + DMSO SM 1:100
U251 + PAV-0805 SM 1:100
ALDH1A1
KAP1
PDI
YWHAZ
